# eIFiso4G editing confers high and broad-spectrum resistance against rice yellow mottle virus

**DOI:** 10.64898/2026.01.30.702252

**Authors:** Florence Auguy, Sophie Chéron, Laurence Dossou, Agnès Pinel-Galzi, Emilie Thomas, Jamel Aribi, Marcos Giovani Celli, Sébastien Cunnac, Eugénie Hébrard, Laurence Albar

## Abstract

Rice yellow mottle virus (RYMV) is one of the most devastating viral pathogens affecting rice cultivation in Africa. The eukaryotic translation initiation factor 4G isoform eIFiso4G1 plays a pivotal role in rice susceptibility to the virus. Naturally occuring resistance alleles impair infection but are predominantly found in the African cultivated species *Oryza glaberrima*, whose use in breeding programs is limited by interspecific sterility barriers with the widely grown *O. sativa*.

Here, we used CRISPR-Cas9 to generate knockout (KO) mutations in eIFiso4G1 as well as insertion-deletion (Indel) variants within the region involved in interaction with the viral protein VPg. KO mutations conferred high resistance to RYMV. No resistance-breaking events were observed, suggesting that this resistance may be more durable than that conferred by naturally occuring alleles. However, complete gene KO slightly affected plant growth. Lines carrying Indel mutations in the VPg-interacting region displayed variable resistance levels, with some variants conferring high resistance without compromising plant development. Structural modeling of the eIFiso4G1 variants and their complexes with VPg provided mechanistic insights into how specific Indels modulate the resistance phenotype.

## Introduction

Viral diseases are a major constraint to crop production, often causing substantial yield losses (Tatineni and Hein, 2023). Some of them, especially those affecting staple food crops, may even threaten food security. Chemical control methods are generally ineffective against virus epidemics, especially in case of mechanical transmission, and have proven harmful to both the environment and consumers. While prophylactic measures and cultural practices remain essential components of integrated management strategies, they are often insufficient on their own. As a result, developing virus-resistant crop varieties is widely regarded as the most effective strategy to mitigate virus-induced yield losses.

Most plant resistances to pathogens, including viruses, rely on dominant resistance genes, such as nucleotide-binding leucine-rich repeat genes, which mediate the recognition of pathogen-derived molecules and trigger immune responses (for review, see Contreras et al., 2023). However, recessive resistance gene**s** are more frequently observed against viruses than against cellular pathogens such as fungi or bacteria (Kourelis and van der Hoorn 2018). Indeed, due to their compact genomes, viruses heavily depend on host cellular factors for their life cycle—factors that can thus act as host susceptibility determinants. The most well-documented examples include the eukaryotic translation initiation factor 4E (eIF4E) and its isoform, which are recruited by several viruses of the *Potyviridae* family to infect both Poaceae and dicotyledonous plant species (Sanfaçon, 2015). Other factors associated with recessive resistance include the translation initiation factors OseIf4G and OseIFiso4G1, the nucleoporin OsCPR5-1, the protein disulfide isomerase-like HvPDIL5.1 or the phosphoglycerate kinase AtcPGK2 (Albar et al., 2006; Lee et al., 2010; Ouibrahim et al., 2014; Yang et al., 2014; Pidon et al., 2020).

For some susceptibility factors, such as OsCPR5.1 or HvPDIL5.1, resistance is conferred by loss-of-function (KO) mutations, making these genes straightforward targets for resistance engineering using CRISPR-Cas9 genome editing technology (Cheng et al., 2022; Arra et al., 2024). For other factors, including the translation initiation factors, most naturally occurring resistance alleles are characterized by point mutations. These mutations do not affect the protein’s native function for the plant, but disrupt its interaction with the viral partner protein, preventing the virus from recruiting the host factor to complete its cycle. Generating KO mutations can also result in a resistance phenotype but it can sometimes lead to developmental defects, such as reduced viability in severe cases, or unexpected consequences like decreased resistance spectrum or durability(Piron et al., 2010; Mazier et al., 2011; Macovei et al., 2018; Kuroiwa et al., 2021; Pechar et al., 2022). In these cases, a more sophisticated approach is to select alleles that retain the protein’s native function for the plant while specifically impairing its interaction with the virus. Such edited alleles could offer a more sustainable resistance strategy and avoid the fitness costs associated with KO mutations (Bastet et al., 2017).

Rice yellow mottle virus (RYMV, *Sobemovirus RYMV*) represents a significant threat to rice cultivation in Africa, where it is endemic. Savary et al. (2019) estimated yield losses at about 4.3% in Sub-Saharan Africa, making yellow mottle one of the most serious diseases of rice in this region. Depending on varietal susceptibility and growing conditions, yield losses can exceed 50%, leading to significant consequences for smallholder farmers who rely heavily on rice for their livelihood (Kouassi et al., 2005). Previous research has identified three major resistance genes : *RYMV1* and *RYMV2*, which encode susceptibility factors—the translation initiation factor OseIFiso4G1 and the nucleoporin OsCPR5.1, respectively (Albar et al., 2006; Orjuela et al., 2013)—and *RYMV3*, which encodes a dominant immune receptor (Bonamy et al., 2023). While resistance sources are mainly found in the African rice species *Oryza glaberrima*, their transfer to the widely cultivated *O. sativa* is hindered by interspecific sterility barriers (Pidon et al., 2020). Furthermore, the identification of RYMV variants capable of overcoming natural resistance alleles highlight the need for innovative approaches (Pinel-Galzi et al., 2016; Hébrard et al., 2018; Bonamy et al., 2023). Gene editing offers a promising solution, enabling the development of resistant *O. sativa* varieties and the generation of potentially more durable resistance alleles. Recent work has already demonstrated the feasibility of this approach for *RYMV2* (Arra et al., 2024). In this study, we used CRISPR/Cas9 to introduce KO mutations and deletions in *OseIFiso4G1*, aiming to disrupt the virus’s ability to hijack the host’s translational machinery. Additionally, we targeted an *RYMV1* homologous gene within the rice genome to investigate its potential role in conferring resistance. Through phenotypic evaluation of the edited lines, we assessed their resistance to RYMV and the impact of the mutations on developmental traits. Our findings contribute to a better understanding of the interactions between RYMV and the rice eIFiso4G factors, and may support the development of new strategies for engineering RYMV resistance through genome editing.

## Material and methods

### Generation of edited plants

CRISPR/Cas9 editing was performed on the *Oryza sativa* cv. Kitaake, a cultivar susceptible to RYMV, targeting the genes *OseIFiso4G1* and *OseIFiso4G2* (corresponding to gene models LOC_Os04g42140 and LOC_Os02g39840 in the Nipponbare reference genome; Kawahara et al., 2013).

For *OseIFiso4G1*, three guide RNAs (gRNA-1.1, gRNA-1.2, and gRNA-1.3) were designed to target the sixth exon, with the aim of inducing KO mutations or small in-frame indels within a 46-bp region associated with RYMV resistance (Fig. 1; Pidon et al., 2020). Each gRNA was inserted into a cassette containing the rice U6.1p promoter, the sgRNA scaffold, and attL1/attL2 recombination sites. For *OseIFiso4G2*, two gRNAs (gRNA-2.1 and gRNA-2.2), located a few dozen base pairs apart, were designed to target the fourth exon (Fig. 1), with the goal of inducing large deletions between the two cut sites. A single cassette containing both gRNAs was synthesized, with U6.1p and U6.2p promoters driving expression of gRNA-2.1 and gRNA-2.2, respectively, along with the sgRNA scaffold and attL1/attL2 sites. All four cassettes were subcloned into the modified binary vector pH-Ubi-Cas9-7 (Fayos et al., 2020; Miao et al., 2013) using the Gateway LR Clonase II reaction (Invitrogen), resulting in the constructs p-iso4G1.1, p-iso4G1.2, p-iso4G1.3, and p-iso4G2. These constructs were introduced into *Agrobacterium tumefaciens* strain EHA105 by electroporation, and rice transformation was carried out via *Agrobacterium*-mediated transformation, following protocols described by Blanvillain-Baufumé et al. (2017) and Sallaud et al. (2003).

**Figure 1:**
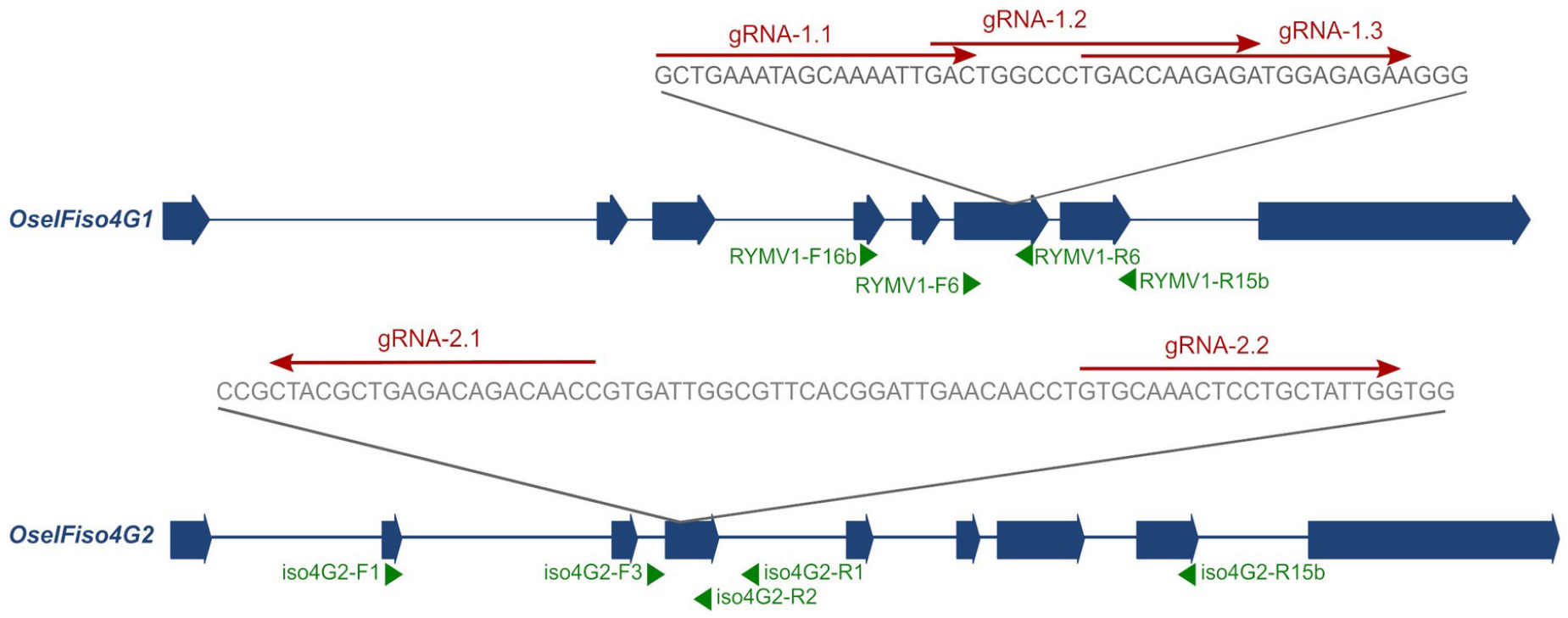
Position of the gRNA designed on *OseIFiso4G1* and *OseIFiso4G2*. Exons and introns are represented by large blue arrows and thin blue traits, respectively; gRNA are represented by red arrows above the sequences; the positions of the primers used in this study are represented by green arrows. Primer iso4G1-R15b is actually straddling exons 7 and 8.

### Genotyping and selection of edited lines

T0 plants were screened for both the presence of T-DNA insertions and mutations in the target genes. PCR amplifications targeting the hygromycin resistance gene and the Cas9 gene were performed using the primer pairs HygR-F (5’-CTCGGAGGGCGAAGAATCTC-3’)/HygR-R (5’-GCTCCAGTCAATGACCGCTG-3’) and Cas9-F (5’-TCACCTCCTTGTAGCCCTTG-3’)/Cas9-R (5’-ACGGCGAGATTAGGAAGAGG-3’), respectively. Mutations in the target genes were analyzed by Sanger sequencing. For *OseIFiso4G1*, amplification was performed with primers RYMV1-F16b (5’-CCTTGGTCAGCTAGAAGAGGCA-3’)/RYMV1-R6 (5’-AGTAGCTCACCAATTAGACGGA-3’), and sequencing was carried out using primer RYMV1-F6 (5’-GAGCCCACCTTCTGTCCGATG-3’). For *OseIFiso4G2*, amplification was done using iso4G2-F1 (5’-AGACTGGTGCAGTTTCACGT-3’)/iso4G2-R1 (5’-AAGCGTATATTGCCCCCTCG-3’), and sequencing with primer iso4G2-R2 (5’-TTTGCACTGAACTGGGAGCT-3’). PCR reactions were carried out either directly on leaf extracts (approx. 1 cm^2^ of leaf tissue ground in 200 µL of water) using the Terra PCR Direct Polymerase Kit (Takara), or on genomic DNA extracted as described by Edwards et al. (1991), using GoTaq G2 Polymerase (Promega), following the respective manufacturers’ protocols. Sequencing was outsourced to Genewiz. Superimposed chromatograms from heterozygous samples were interpreted using DSDecodeM (Liu et al., 2015) and results were manually validated.

T1 and T2 lines were similarly screened by PCR for the presence of the hygromycin resistance gene or the Cas9 gene, and for mutations in the target genes. Indels larger than 30 bp were detected by size polymorphism on agarose gels, while smaller indels were confirmed by Sanger sequencing.

### cDNA analysis

RNA was extracted from the leaves of 2-week-old uninoculated plants using TriReagent (Euromedex) according to the manufacturer’s instructions. The RNA samples were treated with RQ1 RNase-free DNase (Promega) and quantified using a Nanodrop spectrophotometer. Retrotranscription was performed on 2 µg of RNA using 0.5 µg of oligo-dT primer and M-MLV retrotranscriptase (Promega).

To confirm mutations in the transcripts, cDNA amplification was performed using GoTaq G2 Polymerase (Promega), with the primer pairs RYMV1-F16b (5’-CCTTGGTCAGCTAGAAGAGGCA-3’)/RYMV1-R15b (5’-CTCTTCACGTCGAGGCACCCA-3’) for *OseIFiso4G1* and iso4G2-F3 (5’-CGGGTTCGTTACACTAGGGA-3’)/iso4G2-R15b (5’-CTCTTCACGTCGAGGCACCCA-3’) for *OseIFiso4G2*. The amplified products were sent for Sanger sequencing to Genewiz. Sequencing of *OseIFiso4G1* was performed using primer RYMV1-F6, and sequencing of *OseIFiso4G2* was carried out with primer iso4G2-F3.

The expression level of *OseIFiso4G1* was analyzed by RT-qPCR using the Takyon No ROX SYBR 2X MasterMix blue dTTP (Eurogentec) and the primer pair iso4G1-qPCR-F (5’-AGTTTCCGCCCATGGTTTTC-3’) and iso4G1-qPCR-R (5’-TGGTGAAGTTGTTGCACGAC-3’). Samples consist in a pool of leaves from three or four plants. Actin was used as the reference gene and amplified with the primers actin-qPCR-F (5’-GCGTGGACAAAGTTTTCAACCG-3’) and actin-qPCR-R (5’-TCTGGTACCCTCATCAGGCATC-3’).

### Plant culture and evaluation of developmental traits

Rice plants were grown in greenhouse with a day/night temperature of 28°C/22°C and 70% humidity. Natural daylight was supplemented with lamplight for a 14-hour light/24-h photoperiod. Plants were grown in 2-liter pots for multiplication.

Four independent experiments were conducted to assess the impact of mutations in *OseIFiso4G1* and *OseIFiso4G2* on plant development. Three to six plants per genotype were grown in pots (experiments 5 and 8) or in 230 ml plug trays (experiments 6 and 7). Plants height at ripening stage was measured, and in some assays, the number of reproductive tillers and grain weight were also recorded.

### Resistance and resistance-breaking evaluation

For resistance evaluation, up to 12 plants per line and per experiment were sown in plugs or culture trays. Resistance to the RYMV isolate BF1 from Burkina Faso (strain S2), previously used for RYMV1 mapping and cloning (Albar et al., 2006), was assessed in three independent experiments. In addition, a single experiment was conducted for each of the following isolates: CIa (Côte d’Ivoire, strain S2; Brugidou et al., 1995), Mg1 (Madagascar, strain S4; Pinel et al., 2000), Ng106 (Niger, strain S1; Traoré et al., 2010), and Tz11 (Tanzania, strain S1; Abubakar et al., 2003). Mechanical inoculation was performed two weeks after sowing, following the protocol described by Pinel-Galzi et al. (2018). The cultivars *Oryza sativa* cv. Kitaake and IR64 were included as susceptible controls, while near-isogenic IR64 lines carrying the *rymv1-2* or *rymv1-3* alleles served as resistant controls. Symptoms were first recorded between 11 and 14 days after inoculation (DAI), depending on the isolate and the experiment, and a second evaluation was carried out between 17 and 21 DAI. A symptom severity scale ranging from 1 to 9 was used, where scores of 1 and 2 indicated no visible or ambiguous symptoms, respectively, and a score of 9 corresponded to orange discoloration or leaf necrosis (Pinel-Galzi et al., 2018). Samples from the last emerged leaves were collected between 13 and 15 DAI for ELISA testing. Detection of viral accumulation was conducted using direct double-antibody sandwich (DAS) ELISA, as described by Pinel-Galzi et al. (2018), using a polyclonal antiserum raised against a Madagascan RYMV isolate (N’Guessan et al., 2000).

To assess resistance-breaking, up to 96 plants per line and per isolate were tested. Three isolates were used: Ng106, Ng119 (Niger, strain S1; Pidon et al., 2017), and Tg247 (Togo, strain S1; Pinel-Galzi et al., 2016). Inoculation was performed two weeks after sowing, and leaf samples were collected at 64 DAI for ELISA testing. Resistant controls consisted of *O. sativa* cv. Gigante (carrying the *rymv1-2* allele) and *O. glaberrima* cv. Tog5681 (carrying the *rymv1-3* allele). Inoculation efficiency was assessed using five plants each of the susceptible IR64 and Kitaake varieties.

### Statistical analysis

Statistical analyses were performed using the free software R (http://www.r-project.org/). For expression level, resistance levels and plant development traits, Kruskal-Wallis non-parametric tests were conducted for each independent experiment to identify significant differences between lines (p<0,05). Multiple comparisons were performed using the Dunn post-hoc test, and p-values were adjusted using the FDR method (package rstatix). For the evaluation of final plant height, the four experiments were analyzed together using a linear mixed model and Tukey’s post-hoc tests (package emmeans). For resistance-breaking, Fisher’s exact tests (package RVAideMemoire) were performed to identify significant differences between lines and for multiple comparisons.

### Structural modeling of the MIFiso4G/VPg complexes

The central domain of OseIFiso4G1 WT and edited proteins were modeled alone and in interaction with BF1 VPg using AlphaFold v3.3 (Jumper et al., 2021) with defaut parameters, a deep-learning-based modeling tool. Scores of predicted local distance difference test (plDDT), predicted aligned error (PAE), predicted template modelling (pTM) and interface predicted modelling (ipTM) were checked and compared as suggested in the prediction tool. The model of WT MIF4G was superimposed onto the homologous structure present in the Protein DataBank and pairwise root mean square deviations were calculated using PyMol (The PyMOL Molecular Graphics System, Version 3.0 Schrödinger, LLC). The sequence identity between 1HU3 and MIFiso4G1-1 IR64 was calculated after alignment with Ident and Sim (Stothard, 2000). Amino acids specific to the two clades were visualized on the models using PyMol. Predicted interfaces of the five models obtained for each complex were analyzed using service PISA at the European Bioinformatics Institute (http://www.ebi.ac.uk/pdbe/prot_int/pistart.html) (Krissinel and Henrick, 2007).

## Results

### Development and molecular characterization of lines edited in *OseIFiso4G1* and its paralog

The *OseIFiso4G1* gene, implicated in rice susceptibility to RYMV, was edited in *O. sativa* cv. Kitaake to generate RYMV-resistant lines. Gene editing was performed using three distinct gRNAs targeting a 46 bp region within the central domain of the gene, known to interact with the viral VPg. Among the 90 T0 transgenic lines analyzed, 66 were edited in *OseIFiso4G1*, with approximately half being bi-allelic mutants (Sup. Table 1). Seventeen independent T0 with mutations resulting in premature stop codons and eleven with in-frame indels were identified, while mutations in the other T0 remained unresolved due to complex chromatograms. T1 and T2 plants were selected based on the absence of the transgene and the presence of homozygous mutations in *OseIFiso4G1*. Ultimately, three independent lines carrying premature stop codons and seven harboring in-frame indels were chosen for detailed phenotypic characterization. Partial sequencing of the transcript confirmed the expected mutations at the cDNA level and indicated that these mutations did not disrupt the downstream splicing site.

Lines iso4G1-KO3, iso4G1-KO8, iso4G1-KO11 are characterized by premature stop codon, such that less than 50% of the protein sequence was retained (Fig. 2a, Fig. 3a, Sup. Data 1). The eIF4B and one eIF4A interaction domain were missing, while the PABP and the second eIF4A interaction domains were only partially conserved (Gallie, 2016), likely resulting in severe impairment of protein function. These lines are thus considered KO mutants and will be referred to as OseIFiso4G1 KO lines. Lines iso4G1-del1, iso4G1-del2, iso4G1-del3, iso4G1-del4, iso4G1-del6, iso4G1-del7, iso4G1-del9 feature single amino acid substitutions and/or deletions of 1 to 8 amino acids within the conserved central MIF4G domain (Fig. 2a, Fig. 3a, Sup. Data 1). RT-qPCR analysis revealed no significant differences of *OseIFiso4G1* expression levels associated to the deletions (Sup. Fig. 1).

**Figure 2:**
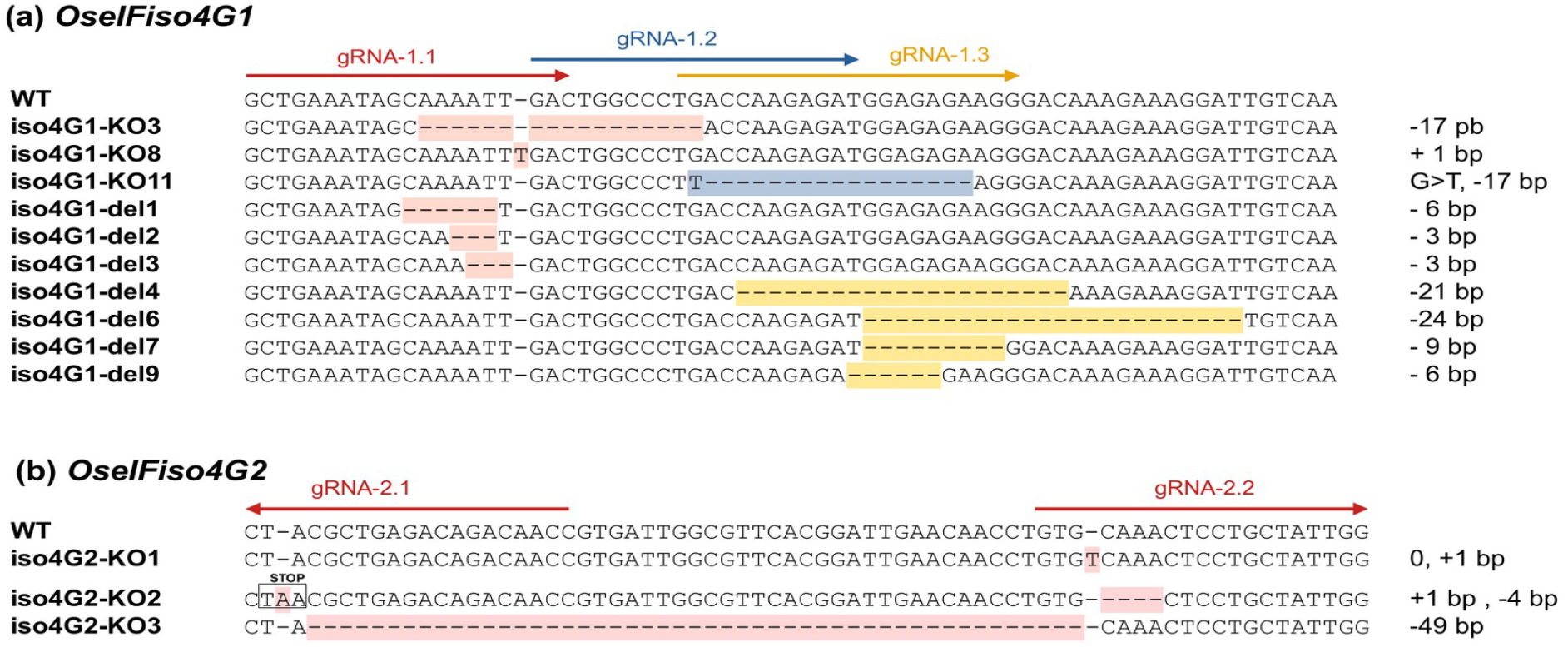
Mutations introduced in the 6th exon of *OseIFiso4G1* (a) and in the 4th exon of *OseIFiso4G2* (b) using CRISPR-Cas9 technology. gRNA are represented by colored arrows. The names of the lines are indicated on the left side of the sequences. The mutations in *OseIFiso4G1* are highlighted with different colors depending on the construct with which they have been obtained. The size of the insertion or deletions, or eventually the substitutions, compared to the Kitaake wild-type sequences are indicated on the right side of the sequences.

**Figure 3:**
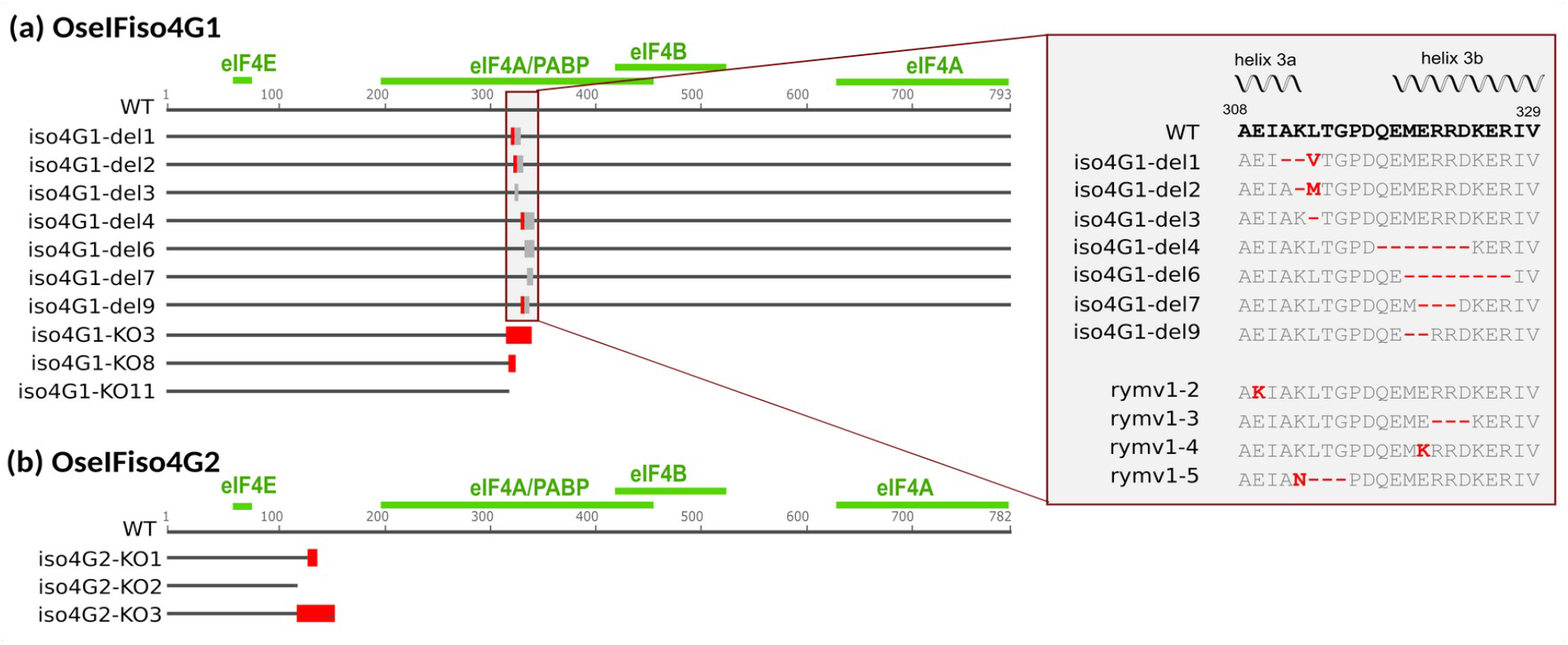
Impact of the mutations introduced in the genes encoding OseIFiso4G1 (a) and OseIFiso4G2 (b) on the resulting proteins. Lines’ names are indicated on the left. The conserved parts of the sequence are represented by a dark grey line, deletions by light grey boxes and substitutions by red boxes. Domain of interactions with other proteins of the translation initiation complex, as identified in Gallie (2016), are represented as green lines above the WT sequences. For lines with in-frame mutations, the amino acid sequences in the mutated region are indicated on the right side of the figure and compared to the WT sequence, corresponding to a susceptibility allele, and to the natural resistance alleles rymv1-2, rymv1-3, rymv1-4 and rymv1-5. The position of helix structures are indicated above the sequence.

In parallel, the *OseIFiso4G2* gene was edited using gRNAs targeting the fourth exon. Due to technical issues in transgenic calli selection, only four transgenic plants were obtained. Two of these were edited on *OseIFiso4G2*: one with a homozygous mutation and another with two different mutations. From these two independent T0 plants, three lines homozygous for distinct mutations on *OseIFiso4G2* were selected for further analysis. Lines iso4G2-KO1 and iso4G2-KO2 are characterized by small insertions or deletions (<4bp) in the target regions, while line iso4G2-KO3 features a 49 bp deletion between the two target sequences (Fig. 2b, Fig. 3b, Sup. Data 1). These mutations result in premature stop codons that theoretically preserved no more than 15% of the protein sequence, with the interaction domains for eIF4A, eIF4B, eIF4E, and PABP completely absent (Gallie, 2016). These lines are therefore considered KO mutants and will be referred to as OseIFiso4G2 KO lines.

In an effort to develop double mutants, five independent crosses were made between an OseIFiso4G1 KO line and an OseIFiso4G2 KO line. Resulting F2 progenies were genotyped for mutations in *OseIFiso4G1* and *OseIFiso4G2*. Screening of 434 F2 plants did not identify any double mutants (Sup. Table 2), suggesting that double mutants are not viable. Additionally, an excess of plants with WT genotype (42% instead of the expected 25%) and a deficit of plants with *OseIFiso4G1* KO mutations (4,8% instead of the expected 25%) were observed. Similar results, albeit to a lesser extent, were also observed for *OseIFiso4G2* (32% WT and 17,8% KO instead of the expected 25%). These results were consistent between the five tested progenies. To confirm these findings, a F3 progeny fixed for a KO mutation in *OseIFiso4G1* and segregating for *OseIFiso4G2* were screened. The observed segregation (77 WT, 60 heterozygous, 0 KO) confirmed the absence of viable double mutants. The grain filling rate in F2 plants homozygous for a KO mutation in one *OseIFiso4G* copy and heterozygous for the other one suggests that female sterility or grain abortion alone cannot fully account for the absence of double mutants. Similarly, the germination rate of their F3 progenies suggests that a germination defect alone is also insufficient to explain this absence (Sup. Table 3). The absence of double mutants could thus be explained by impaired pollen viability or by a combination of factors.

### Phenotyping of edited lines for RYMV resistance

Homozygous edited lines were challenged with the BF1 isolate of RYMV. Symptom development was monitored at 11 and 18 days after inoculation (DAI) across three independent experiments. In two of these experiments, leaf samples were collected at 13 DAI for ELISA-based detection of viral accumulation. Despite some variations in disease severity between experiments, all three led to the same conclusions. Lines isoG1-KO3, isoG1-KO8, isoG1-KO11, characterized by frameshift mutations and premature stop codons in *OseIFiso4G1* showed high resistance, with no clear symptoms 18 DAI and no virus detected using ELISA (Fig. 4a-b, Sup. Fig. 2). These results clearly demonstrate that KO mutations in *OseIFiso4G1* confer resistance to RYMV. Lines iso4G1-del4 and iso4G1-del6, featuring seven and eight amino acid deletions in the MIF4G central domain of *OseIFiso4G1*, respectively, exhibited resistance levels similar to those of OseIFiso4G1 KO lines. Line iso4G1-del7, characterized by a three amino acid deletion, developed symptoms at 18 DAI and allowed virus multiplication, but was significantly less susceptible than the Kitaake WT in most cases (Fig. 4a-b, Sup. Fig. 2). The four other lines with in frame mutations (iso4G1-del1, iso4G1-del2, iso4G1-del3, iso4G1-del9) were generally not significantly different from the Kitaake WT. The three lines with KO mutations in *OseIFiso4G2* showed virus content and symptom severity that were not significantly different than those observed in the Kitaake WT, suggesting that *OseIFiso4G2* is not involved in RYMV resistance.

**Figure 4:**
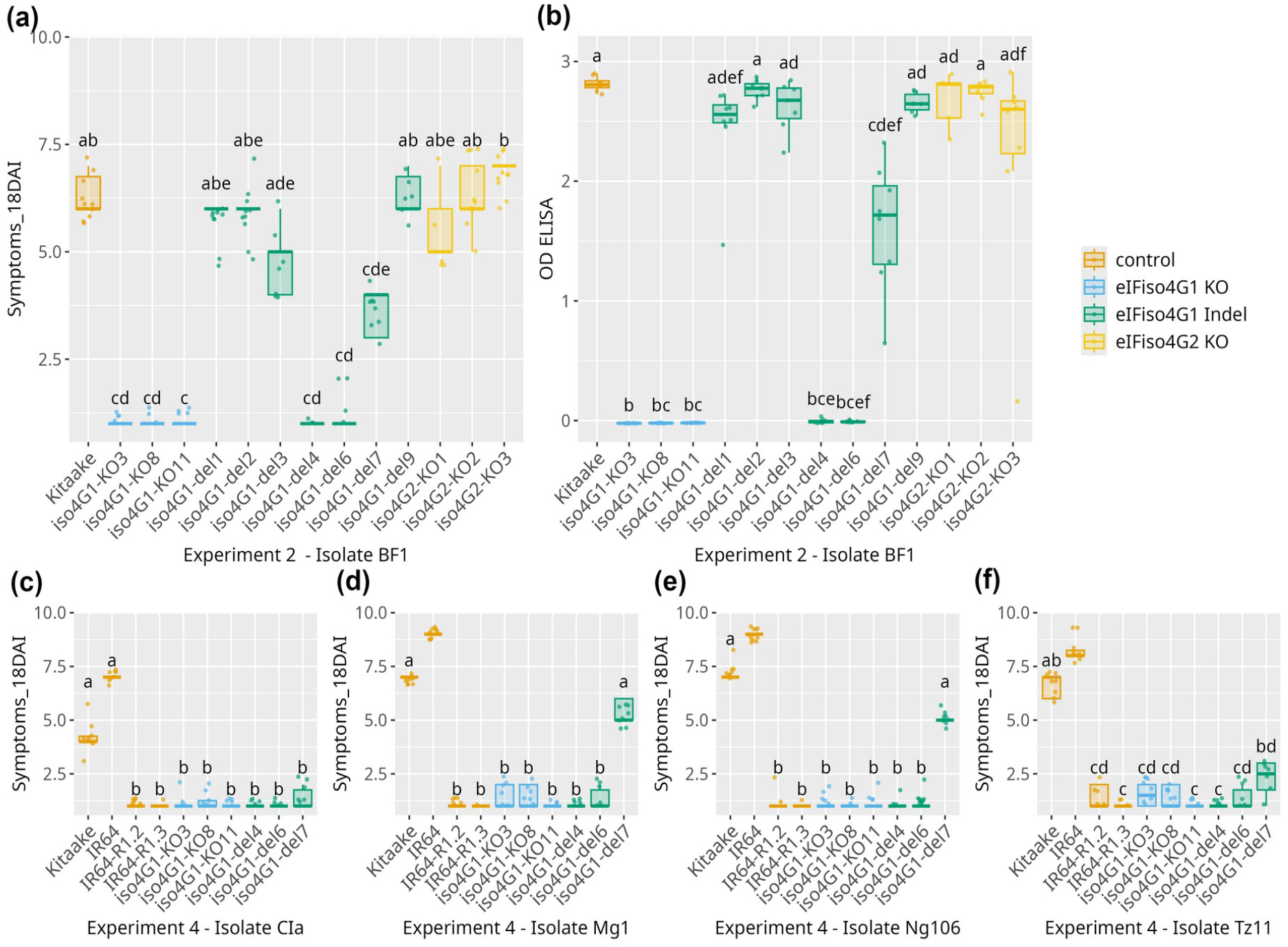
Effect of edition in OseIFiso4G1 and OseIFiso4G2 on resistance against RYMV. Resistance was evaluated based on symptoms and DAS-ELISA. Here are represented symptoms 18 DAI after inoculation of BF1 isolate (a), OD (DAS-ELISA) on samples collected 13 DAI after BF1 inoculation (b), and symptoms 18 DAI after inoculation of isolates CIa (c), Mg1 (d), Ng106 (e), Tz11 (f). Results of a single experiment among 4 performed with BF1 are represented. Additional results are illustrated in Sup. Fig. 2 and Sup. Fig. 3. The wild type Kitaake and IR64 varieties were used as susceptible controls. IR64 variety introgressed with resistance alleles *rymv1*.*2* (IR64_R1.2) or *rymv1*.*3* (IR64_R1.3) were used as resistant controls. The lower and upper hinges correspond to the first and third quartiles; the upper vs. lower whisker extend from the hinge to the largets vs. smallest value no further than 1,5 x interquartile from the hinge. Different letters above the box-plots represent significant differences based on a Kruskall-Wallis test, followed by a Dunn post-hoc test (p-value <0,05 after fdr correction).

Lines showing high or partial resistance to the BF1 isolate were also tested for resistance to four additional isolates representing the diversity of RYMV: CIa, Ng106, Mg1 and Tz11. Lines with KO mutations (iso4G1-KO3, -KO8, -KO11) and lines iso4G1-del4 and iso4G1-del6 were highly resistant to all four isolates, as previously observed for isolate BF1 (Fig. 4c-f, Sup. Fig. 3). All four isolates were able to infect the iso4G1-del7 line, as indicated by positive ELISA OD values. However, symptoms were consistently milder than in the Kitaake WT, although not always significantly so. These results suggest that the iso4G1-del7 line exhibits partial resistance, while variation in symptom severity among isolates may reflect differences in aggressiveness or isolate

× genotype interactions, or a combination of both factors.

### Ability of RYMV to overcome KO mutations in *OseIFiso4G1*

We assessed the ability of RYMV to overcome the resistance in the edited lines iso4G1-KO3, iso4G1-KO8, iso4G1-KO11 as well as in the resistant controls *O. sativa* cv. Gigante and *O. glaberrima* cv. Tog5681, which carry the *rymv1*.*2* and *rymv1*.*3* resistance alleles, respectively. Isolates Ng106, Ng119 or Tg247 were selected for their strong capacity to overcome the *RYMV1* resistance gene (Hébrard et al., 2018). Eight weeks post-inoculation, all three isolates successfully overcame Tog5681 resistance with infection rates ranging from 22 to 86% (Fig. 5, Sup. Table 4). Ng106 and Ng119 also overcame Gigante with infection rates of 3 and 79%, respectively, whereas no resistance-breaking events were observed with Tg247 on this genotype. In contrast, none of the isolates were able to overcome resistance in the edited lines iso4G1-KO3, -KO8, or -KO11. These results strongly suggest that KO mutations in the susceptibility gene *OseIFiso4G1* are much more difficult to overcome than the naturally occurring point mutations found in *rymv1*.*2* and *rymv1*.*3* alleles.

**Figure 5:**
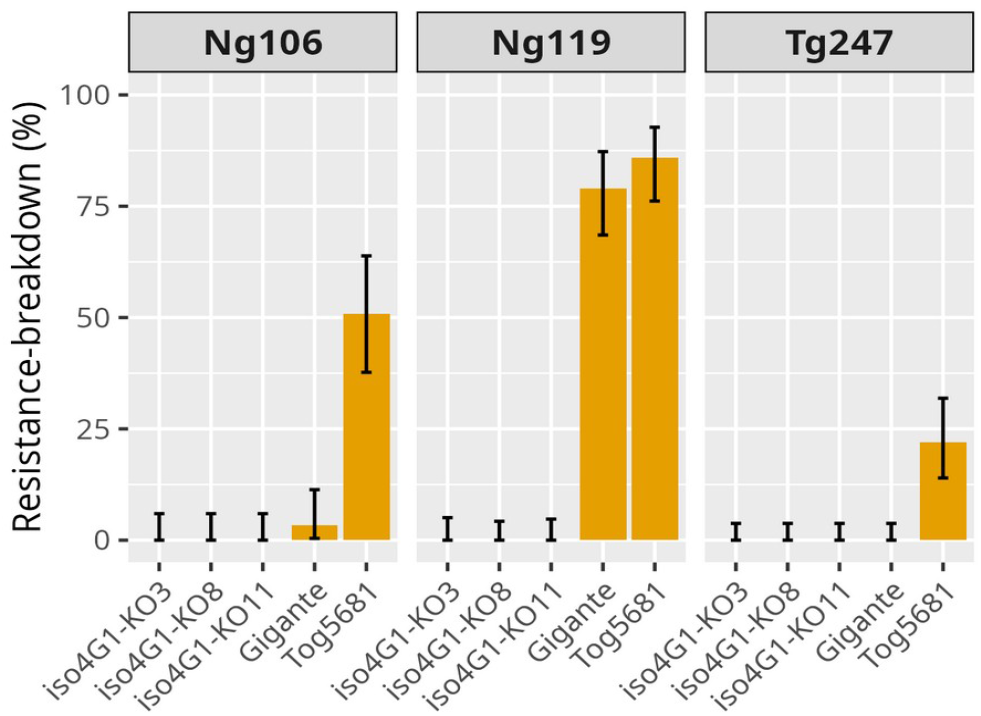
Effect of KO mutations in *OseIFiso4G1* on resistance-breakdown rate. Resistance-breaking rate was measured by the number of plants detected as infected based on a ELISA test performed on leaf samples collected 8 weeks after inoculation.Three different isolates were used. The naturally resistant accessions Gigante and Tog5681, carrying *rymv1-2* and *rymv1-3* resistance alleles respectively, were evaluated as controls. The error bar represents the confidence interval at 95 % calculated using an exact binomial test.

### Effect of mutations in *OseIFiso4G1* and *OseIFiso4G2* on developmental traits

The effect of mutations in *OseIFiso4G1* and *OseIFiso4G2* on developmental traits was evaluated under controlled conditions. Plant height at ripening stage was measured in four independent experiments, each with 3 to 6 plants per genotype. KO mutations in *OseIFiso4G2*, and to a lesser extent in *OseIFiso4G1*, result in a significantly reduced plant height compared to the Kitaake wild type (Fig. 6, Sup. Fig. 4). Lines with in-frame deletions in *OseIFiso4G1* are not significantly different from the control, except for line iso4G1-del9 which is smaller. The number of fertile tillers and seed weight were also measured in, two and one experiment, respectively. No significant differences with the control were found, possibly due to a low statistical power, but the results obtained for seed weight suggested the same trend as for plant height.

**Figure 6:**
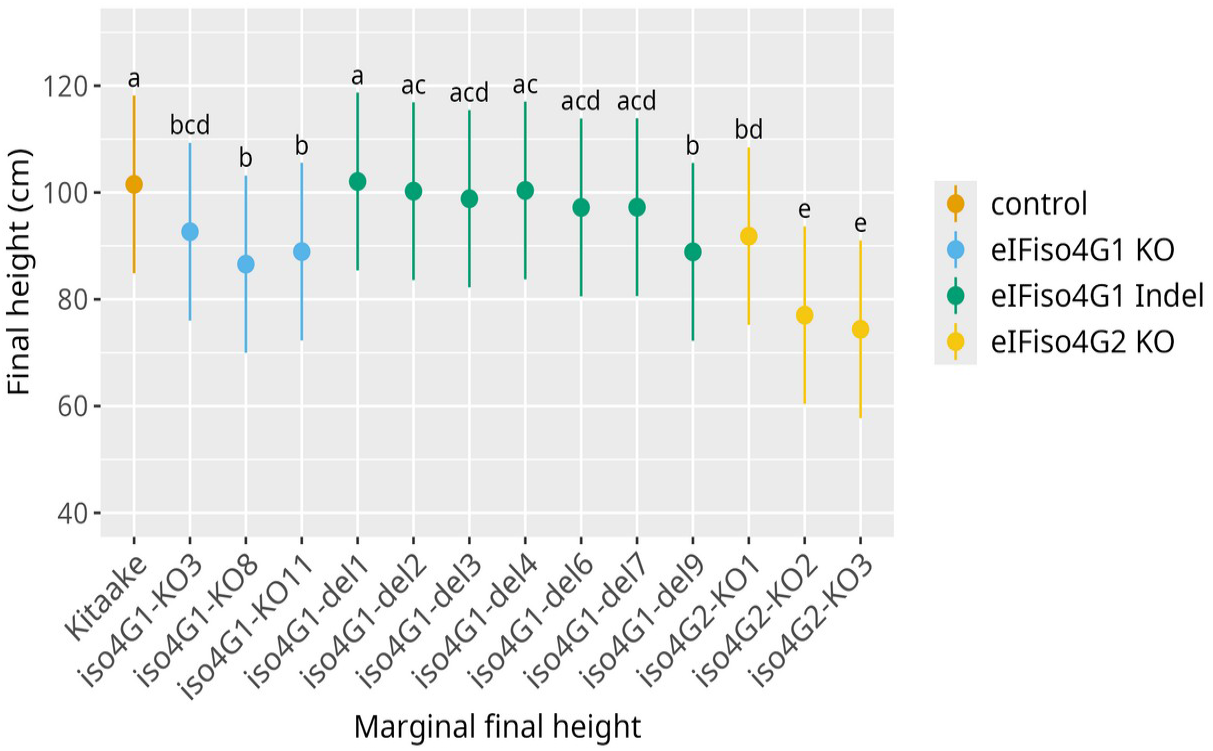
Effect of edition in *OseIFiso4G1* and *OseIFiso4G2* on plant height. Plant height at ripening stage was measured in 4 independent experiments, using 4 to 6 plants per genotype. Here are represented marginal means and confidence intervals at 0,95 % estimated by a linear mixed model on the results of the 4 experiments. Different letters above the box-plots represent significant differences, as estimated by a Tuckey test. Results of individual experiments are illustrated in Sup. Fig. 4.

### In frame mutation impacts on predicted 3D structures

3D structural modeling was performed to assess the potential functional impact of in-frame mutations on OseIFiso4G1 and on its complex with the VPg of RYMV. As a first step, the 3D structure of the central domain (MIF4G, residues 208-435) was predicted for the Kitaake WT protein using AlphaFold v3.3. High predicted local distance difference test scores (plDDT> 70%) and low predicted aligned error (PAE<5%) were observed, indicating highly reliable to reliable model, apart from a local confidence decrease in one loop of the C-terminal region (Fig. 7a, b). The predicted model exhibited strong structural similarity to the human MIF4GII structure resolved by crystallography (PDB 1HU3) with root mean square deviations (RMSD) below 2.5 angstroms. Consistent with previous report (Albar et al., 2006), the two central helices, 3a and 3b, were predicted to form a longer hairpin in rice (Sup. Fig. 5). This hairpin harbors the naturally occurring resistance mutations and is thought to interact with the VPg of RYMV.

**Figure 7:**
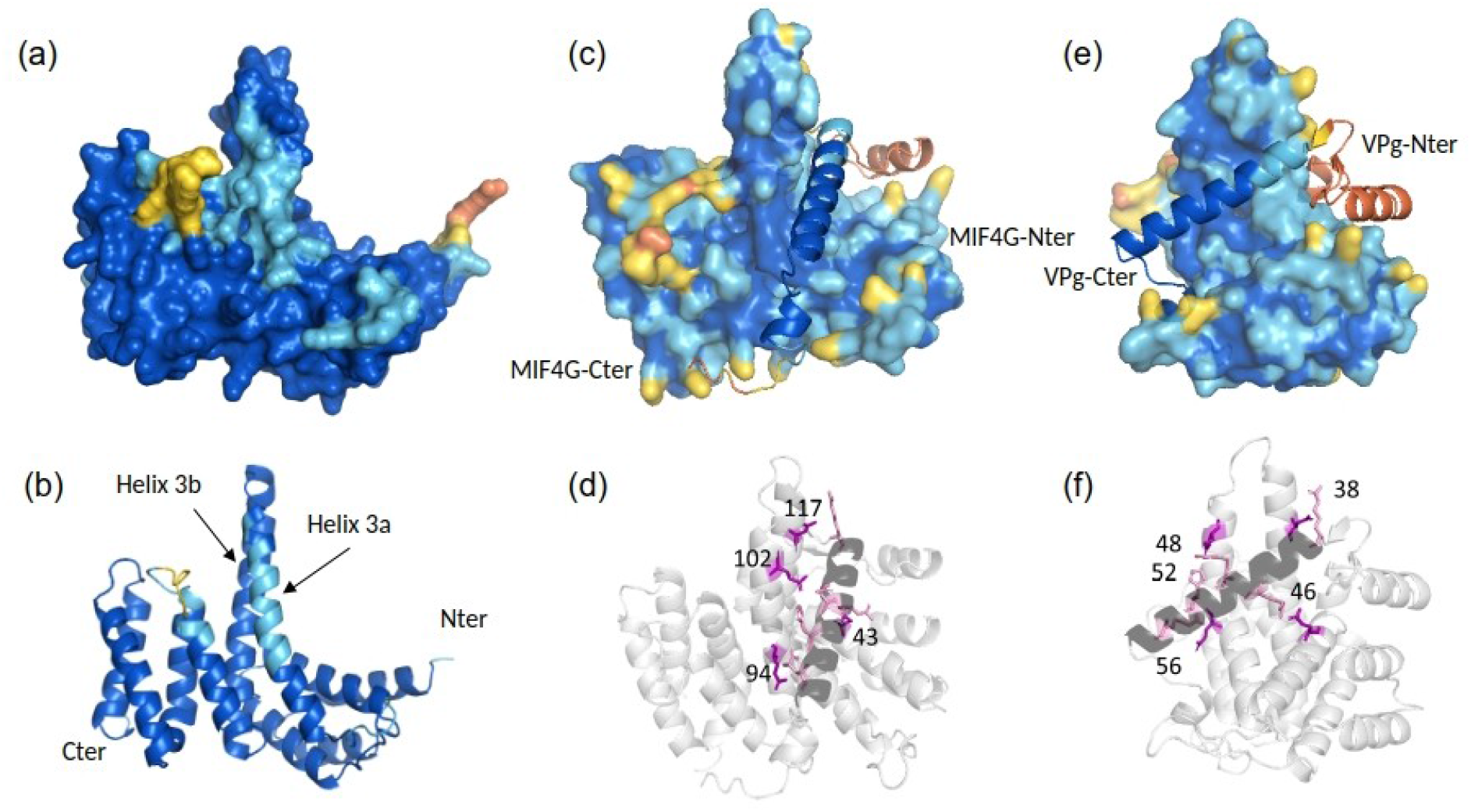
Predicted 3D structure of the rice MIF4G domain (a, b) and of its complex with RYMV VPg (c-f). The models of the MIF4G domain from the Kitaake variety are shown as surface (a, c, e) and cartoon (b, d, f) representations, while the VPg is represented as cartoon in all representations (c-f). The models are displayed in side views (a-d) and face views (e-f). The structures are colored according to per-residue model confidence score (plDDT): dark blue (plDDT>90), high-confidence regions; light blue (plDDT>80), confident regions; yellow (plDDT>60) and orange (plDDT>50), low-confidence regions. The N- and C-terminal extremities of MIF4G (b, c), and VPg (e), and the helices 3a and 3b of MIF4G (b) are indicated. Key residues of MIF4G (numbering on (d)) and VPg (numbering on (f)) involved in the interaction interface are shown on the cartoon representations as purple and pink sticks, respectively (d, f).

Secondly, the MIF4G/VPg complex was modeled to identify the main residues of WT MIF4G required for a compatible interaction. The 3D conformation of the VPg, masked by its intrinsically disordered nature was revealed through its interaction with MIF4G (Hébrard et al., 2009). In complex with MIF4G, the C-terminal half of VPg was predicted to fold mainly as a long central alpha-helix, with very high plDDT scores, while the structure of the N-terminal half of VPg remains unpredictable (Fig. 7c-f). Noteworthy, the 3D prediction of the complex was not impacted by the in silico deletion of the VPg N-terminal domain. Nevertheless, for the remainder of the analyses, MIF4G and its variants were modeled in complex with the entire VPg because indirect effects of this domain cannot be excluded. The complex showed a low PAE score (< 5 Å except for the Nterminal of the VPg) and a high interface predicted template modeling score (ipTM=0,84), allowing inferences of predicted atomic contacts at the interface with high confidence. PISA analysis revealed five main salt bridges between VPg residues at positions 38, 46, 48, 52 and 56, mainly within the central helix, and MIF4G residues at positions 324, 250, 309 (x2) and 301 (numbering according to the full-length OseIFiso4G1 protein), mainly in the 3a/b hairpin (Fig. 7d, f).

In a third step, each edited form of MIF4G was modeled both alone and in complex with VPg. High plDDT scores (> 70%) were obtained, apart from the MIF4G hairpin loop and/or the VPg central helix with three variants (iso4G1-del4, -del6, -del7). All the complexes showed a high interface predicted template modeling score (ipTM>0,8). Mutations in lines iso4G-del1, -del2, -del3 and -del9, which are all susceptible to RYMV, did not drastically affect the overall 3D structure of either the MIF4G domain or the MIF4G/VPg complex (Sup. Fig. 6, Sup. Fig. 7). In contrast, in the resistant lines iso4G1-del4 and -del6, and the partially resistant line -del7, the truncated hairpin likely impacted the interaction interface and the VPg folding, as the size and orientation of the VPg central helix differed from those in the WT MIF4G/VPg complex (Fig. 8, Sup. Fig. 7). PISA analysis revealed a significant reduction of the interaction area for the three variants associated with resistance, and higher solvation energies with iso4G1-del4 and -del7 variants, suggesting lower complex stability (Fig. 9). Moreover, while the total number of hydrogen bonds was comparable accross WT and variant complexes, salt-bridge patterns displayed notable variations (Fig. 9, Sup. Table 5). Several MIF4G residues of iso4G1-del4 variants were engaged in interactions with the same key VPg residues in the same time, which could decrease the strength of the interactions and the stability of the interface. For iso4G1-del6 variant, the VPg residues 38 and 46, within the central helix, were not systematically stabilized by salt bridges with MIF4G residues, likely explaining a reduced interface area, despite a low solvation energy. The MIF4G of iso4G1-del7 variant, carrying a three amino acid deletion and resulting in a less reliable model, showed an intermediary situation.

**Figure 8:**
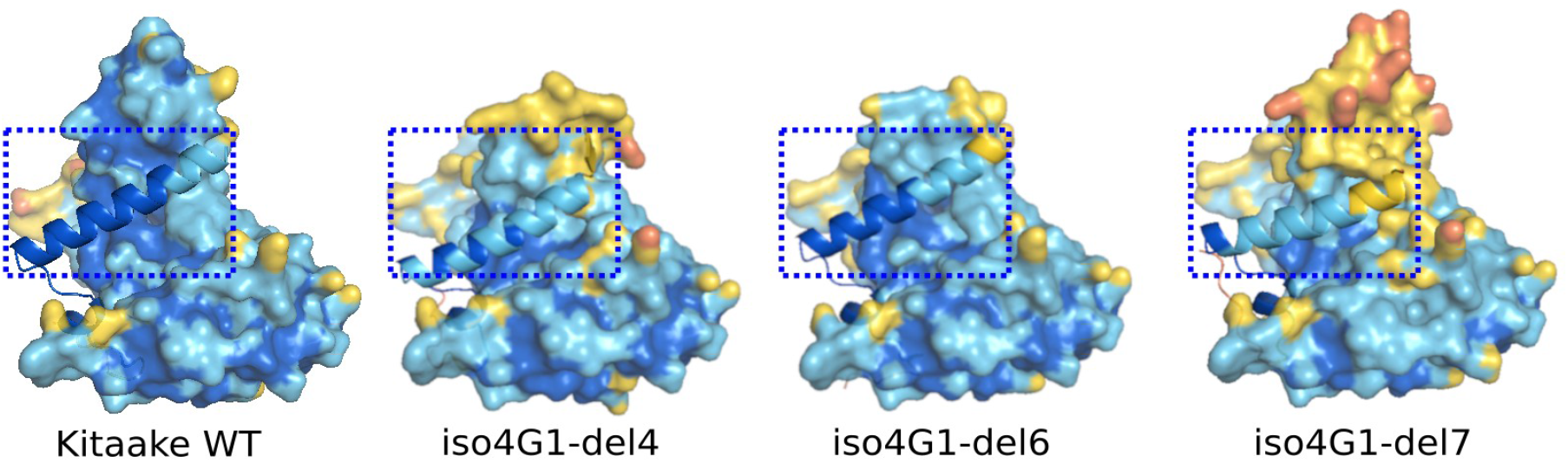
Predicted 3D structure of MIF4G/VPg complex for WT Kitaake and edited variants iso4G1-del4, -del6, -del7. VPg and MIF4G are shown as ribbon and surface representations, respectively. The front view highlights differences in VPg helix length and position among variants. All complexes were superimposed in the same orientation for comparison. The N-terminal domain of VPg was masked for clarity. Structures are colored according to per-residue model confidence score (plDDT), as in figure 6.

**Figure 9:**
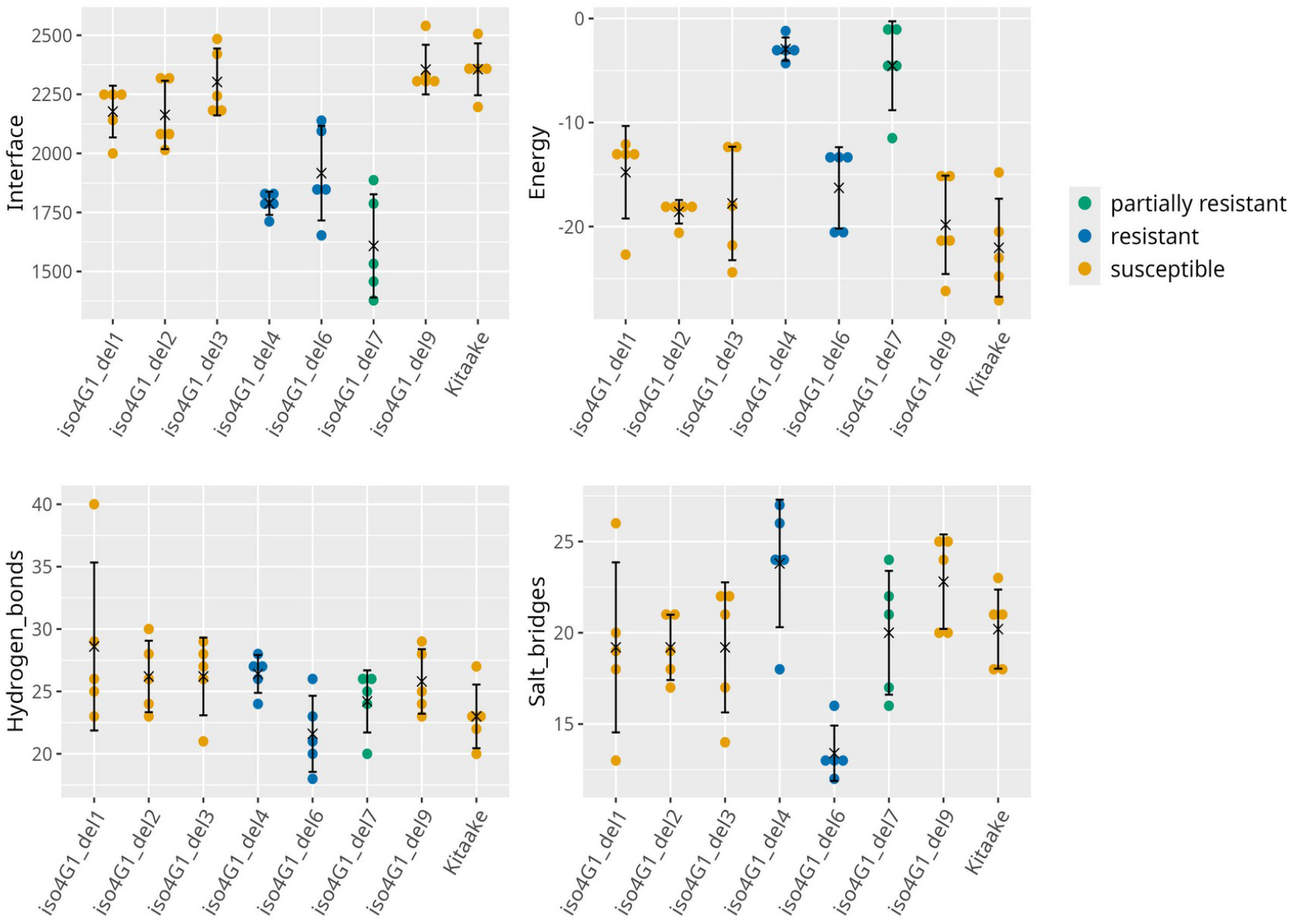
Predicted protein interface analyses using PISA. For each variant, the surface area of the interface expressed in angstroms^2^, the energy of solvation in kcal/mol and the total numbers of hydrogen bonds and salt bridges were counted from each of the 5 models generated by Alphafold. Colours have been assigned to the different lines based on their resistance level, as previously assessed.

## Discussion

In this study, we aimed to generate rice lines resistant to RYMV by editing the *RYMV1* gene, which encodes the translation initiation factor OseIFiso4G1 that plays a pivotal role in rice susceptibility to the virus (Albar et al., 2006). Given that the rice genome harbors two paralogous copies of Os*eIFiso4G*, we analyzed the effects of KO mutations in both *OseIFiso4G1* and *OseIFiso4G2* on viral resistance and plant development (Sup. Table 6). For *OseIFiso4G1*, we additionally produced short deletions in the aim to disrupt viral interaction while maintaining native protein functionality.

We obtained truncated protein variants for both *OseIFiso4G* paralogs. Given the absence or truncation of interaction domains with other translation factors (eIF4A and PABP in OseIFiso4G1; eIF4A, eIF4B, eIF4E, and PABP in OseIFiso4G2), the resulting proteins are regarded as non-functional KO mutants. Although single KO mutants were viable, we observed small but statistically significant reductions in plant height at ripening stage compared with wild type, with a stronger effect of KO in *OseIFiso4G2* than in *OseIFiso4G1*. No significant differences were detected for tillering and grain weight, but the limited number of plants analyzed and the greenhouse culture conditions prevent definitive conclusions.

In contrast, we were unable to recover double KO mutants for *OseIFiso4G1/OseIFiso4G2*, despite screening large numbers of plants and our results suggest that the absence of double mutants might be associated with pollen sterility. This finding indicates that *OseIF4G* cannot complement the simultaneous loss of both *OseIFiso4G* copies. Reciprocally, Macovei et al. (2018) were not able to obtain *OseIF4G* KO mutants, implying that the *OseIFiso4G* genes do not compensate for the absence of *OseIF4G*. Conversely, the viability of single *OseIFiso4G* mutant compared to the non viability for the double mutant points to functional redundancy between the two paralogs for plant development, such that the absence of one gene can be compensated by the other, at least under our experimental conditions. A similar redundancy between *AteIFiso4G* copies has been reported in *Arabidopsis*, despite greater divergence between the two copies than in rice (Gallie, 2016). However, the *Arabidopsis* double *AteIFiso4G* KO mutant, while developmentally affected, remains viable (Lellis et al., 2010), and the single *AteIF4G* KO mutant displays no phenotype (Chen et al., 2014), indicating species-specific differences in the functional relationships among the different translation initiation factor isoforms.

Consistent with the model proposed by Hébrard et al. (2010), who demonstrated that interaction between the RYMV VPg and OseIFiso4G1 is essential for RYMV infection, *OseIFiso4G1* KO mutants showed complete resistance to the virus. No symptoms or viral accumulation were detected, and resistance level matched that conferred by naturally occurring *RYMV1* alleles carrying point mutations that disrupt the VPg-eIFiso4G1 interaction. Importantly, no resistance breakdown was observed in any of the KO lines, even when challenged with the virulent isolate Ng119, which efficiently overcomes all the known resistance alleles of *RYMV1* (Hébrard et al., 2018). These findings suggest that complete loss of OseIFiso4G1 prevents viral adaptation and support the model that *RYMV1* resistance breakdown arises from viral adaptation to modified forms of eIFiso4G1, as proposed by Hébrard et al. (2010), rather than an adaptation to eIFiso4G2. In other plant-virus systems, the situation appears more complex. For instance, in some Solanaceae-potyvirus pathosystems, one eIF4E isoform is preferentially used but viruses can switch to an alternative isoform, when the preferred one is non functional (Gauffier et al., 2016; Moury et al., 2020). In the rice/RYMV pathosystem, Poulicard (2010) reported residual VPg binding to eIFiso4G2 in preliminary yeast double hybrid assays that could have suggested a potential for viral adaptation to this isoform. However our results suggest either strong constraints on viral adaptation to efficiently interact with OseIFiso4G2 or that this isoform is not functionally compatible with the infection process. Nevertheless, we cannot exclude that adaptation to OseIFiso4G2 might occur at a very low frequency.

Beyond the potential for some viruses to exploit alternative susceptibility factors, cross-effects between different translation initiation factor isoforms can lead to unexpected drawbacks in KO mutants. Zafirov et al. (2020) showed that in Arabidopsis, KO in AteIF4E1 confers resistance to clover yellow vein virus (ClYVV, *Potyvirus trifolii*) but increases susceptibility to turnip mosaic virus (TuMV, *Potyvirus rapae*), which preferentially uses the AteIFiso4E isoform. Such cross-effects highlight the potential risks associated with using complete loss-of-function mutations as a resistance strategy. In rice, however, the two known viruses that depend on members of the eIF4G family—RYMV and rice tungro spherical virus (RTSV, *Waikavirus oryzae*)— belong to distinct virus families (Solemoviridae for RYMV and Secoviridae for RTSV), which notably differ in VPg size (9-12 kDa for Solemoviridae vs 2-4 kDa for Secoviridae). This suggests different mode of interactions between VPg and translation initiation factors for RYMV and RTSV, consistent with the role of eIF4G, rather than one of its isoform, in RTSV resistance. Furthermore, these viruses have so far been reported from non-overlapping geographical regions, with RYMV confined to Africa and RTSV to Asia. Consequently, even if KO mutants of *OseIFiso4G1* were to affect RTSV infection, their deployment to control RYMV in Africa would not heighten the risk of Tungro disease—at least under the present distribution patterns of these viruses. Other rice-infecting viruses are present in Africa, notably rice stripe necrosis virus (RSNV, *Benyvirus oryzae*; Bagayoko et al., 2021) and maize streak virus (MSV, *Mastrevirus storeyi*; Fouad et al., 2024). While it remains possible that these viruses hijack plant translation initiation factors, no such mechanism has been reported for viruses within their respective families, which, moreover, lack a VPg.

In addition to generating KO mutants, we produced seven *O. sativa* lines carrying deletions and, in some cases, substitutions within the central domain of OseIFiso4G1. We cannot rule out the possibility that these mutations may affect protein content through transcriptional or translational regulatory mechanisms, even though we did not detect any significant effects on *OseIFiso4G1* expression level. However, given their location in the domain involved in interactions with eIF4A and PABP as well as within the hairpin loop that plays a crucial role in the interaction with the viral VPg, we hypothesized that these mutations could disrupt protein-protein interactions. To explore this, we investigated their impact on structural modeling. Our results suggested that mutations of lines iso4G-del1, -del2, and -del3 did not markedly modify the 3D conformation of the MIF4G domain nor its interaction with VPg. Consistent with the minimal structural impact, these lines exhibited normal growth and remained fully susceptible to RYMV infection. The iso4G1-del9 mutation appeared to cause only minor structural changes to the MIF4G domain and the MIF4G/VPg complex. Consistently, this mutation did not alter susceptibility to RYMV. Nevertheless it resulted in growth defects similar to those observed in the KO lines, suggesting that the interaction of MIF4G with host translation factors eIF4A or PABP might be impaired. In contrast, mutations in lines iso4G1-del4, -del6, and -del7 most strongly disrupted the predicted MIF4G-VPg interface, which correlates with their resistance phenotypes: iso4G1-del4 and -del6 lines exhibited high levels of resistance, while iso4G1-del7 showed partial resistance. Notably, these deletions did not affect plant height, unlike the KO mutants, suggesting that the essential function of OseIFiso4G1 is likely preserved. This finding is somewhat unexpected given the fact that all three mutations affect amino acids conserved across several plant species (Gallie, 2016). Interestingly, the mutations in lines iso4G1-del4, -del6, and -del7 overlap with the three-amino-acid deletion in the *rymv1-3* natural resistance allele, highlighting the critical role of these residues in the interaction with VPg. The mutation in iso4G1-del7, which confers partial resistance, and *rymv1-3*, which confers high resistance, differ by only a single amino acid, demonstrating how a single residue can alter the interaction. However, direct comparison of the predicted 3D structures of our edited variants with *rymv1-3* is complicated by the conformational variability in helix 3a of *rymv1-3*, which varies across models (Sup. Fig. 8).

In summary, this study demonstrates that resistance to RYMV can be achieved through targeted editing of the *OseIFiso4G1* susceptibility gene, confirming that, as previously shown for *RYMV2* (Arra et al., 2024), *RYMV1* gene editing represents a valuable strategy for rice resistance breeding. KO mutations provided strong and potentially durable resistance, while specific in-frame deletions, such as *iso4G1-del4* and *-del6*, conferred high resistance levels without detectable effects on plant growth. Future work will assess the ability of RYMV to overcome the resistance conferred by these deletions. Furthermore, recent advances in protein structure prediction also enabled us to explore the structural consequences of these mutations. Our modeling analyses provide new insights into how specific alterations within the MIF4G domain may influence the stability of the MIF4G-VPg interaction and, consequently, viral susceptibility. These results provide a framework for future structure-guided genome editing approaches aimed at achieving durable resistance to RYMV.

## Acknowledgements

We would like to thank Harold Chrestin, Oscar Main and Pascaline Boyer for technical help, and Sylvain Marthey and Gwenaelle Andre for helpful discussions. A preprint version of this article has been peer-reviewed and recommended by PCI Plants (https://doi.org/10.24072/pci.plants.100070).

## Data, scripts, code, and supplementary information availability

Data are available online: https://doi.org/10.23708/ZWRXNJ (Albar et al., 2026)

