## Supplemental material for "eIFiso4G editing confers high and broad-spectrum resistance against rice yellow mottle virus"

Peer-reviewed & recommended by PCI Plants

### Supplemental material

Sup. Data 1 - Sequences of the eIFiso4G1 and eIFiso4G2 protein in the Kitaake variety and in the different mutant lines.

Sup. Table 1 - Number and genotypes of T0 and T2 plants obtained with the different constructs.

Sup. Table 2 - Segregation of KO and WT alleles in F2 populations derived from crosses between OseIFiso4G1 KO and OseIFiso4G2 KO lines.

Sup Table 3 - Grain filling rate in F2 plants and germination rate in F3 progenies derived from crosses between OseIFiso4G1 KO and OseIFiso4G2 KO lines.

Sup. Table 4 - Resistance-breaking rates on OseIFiso4G1 KO lines.

Sup. Table 5 - Residues involved in key salt-bridges within the VPg central helix–MIF4G hairpin complex.

Sup. Table 6 - Summary of phenotypic traits in the edited lines.

Sup. Fig. 1 - Expression level of OseIFiso4G1 in lines with deletions in the central domain of the gene.

Sup. Fig. 2 - Resistance of edited lines against BF1 isolate of RYMV.

Sup. Fig. 3 - Resistance of edited lines against isolates CIa, Ng106, Mg1 and Tz11.

Sup. Fig. 4 - Morphology of edited lines in controlled conditions.

Sup. Fig. 5 - Comparison between the predicted 3D model of the rice MIF4G domain and the crystallographic structure of human MIF4GII.

Sup. Fig. 6 - MIF4G 3D models of WT Kitaake and edited variants.

Sup. Fig. 7 - Predicted 3D structure of MIF4G/VPg complex for WT Kitaake and edited variants.

Sup. Fig. 8 - Conformational variability in the predicted 3D structures of the MIF4G domain for the *rymv1-3* allele.

**Sup. Data 1- Sequences of the eIFiso4G1 and eIFiso4G2 protein in the Kitaake variety and in the different mutant lines.** Differences with the Kitaake wild type sequence are underlined in red.

**>eIFiso4G1-Kitaake**
MEKDHQPVISLRPGGGGGGPRPGRLFSPAFAAAASGSGDLLRSHVGGASKIGDPNFEVRERVRYTRDQLLELREIVDIPE
AILRINQEIDIELHGEDQIWGRPESDVQVQTQTQAQPHNRYGETDNRDWRARTVQPPAANEEKSWDNIREAKAAHASSGR
QQEQVNRQDQLNHQFASKAQVGPTPALIKAEVPWSARRGNLSEKDRVLKTVKGILNKLTPEKFDLLKGQLMESGITTADI
LKDVISLIFEKAVFEPTFCPMYAQLCSDLNEKLPSFPSEEPGGKEITFKRVLLNNCQEAFEGAESLRAEIAKLTGPDQEM
ERRDKERIVKLRTLGNIRLIGELLKQKMVPEKIVHHIVQELLGSGPDKKACPEEENVEAICQFFNTIGKQLDENPKSRRI
NDTYFIQMKELTTNLQLAPRLRFMVRDVVDLRSNNWVPRREEIKAKTISEIHDEAMKTLGLRPGATGLTRNGRNAPGGPL
SPGGFPMNRPGTGGMMPGMPGTPGMPGSRKMPGMPGLDNDNWEVPRSKSMPRGDSLRNQGPLLNKPSSINKPSSINSRLL
PHGSGALIGKSALLGSGGPPSRPSSLMASLTHTPAQTAPSPKPVSAAPAVVPVTDKAAGSSHEMPAAVQKKTVSLLEEYF
GIRILDEAQQCIEELQCPEYYSEIVKEAINLALDKGPNFIDPLVRLLEHLHAKKIFKTEDLKTGCLLYAALLEDIGIDLP
LAPALFGEVVARLSLSCGLSFEVVEEILKAVEDTYFRKGIFDAVMKTMGGNSSGQAILSSHAVVIDACNKLLK*

**>eIFiso4G1-KO3**
MEKDHQPVISLRPGGGGGGPRPGRLFSPAFAAAASGSGDLLRSHVGGASKIGDPNFEVRERVRYTRDQLLELREIVDIPE
AILRINQEIDIELHGEDQIWGRPESDVQVQTQTQAQPHNRYGETDNRDWRARTVQPPAANEEKSWDNIREAKAAHASSGR
QQEQVNRQDQLNHQFASKAQVGPTPALIKAEVPWSARRGNLSEKDRVLKTVKGILNKLTPEKFDLLKGQLMESGITTADI
LKDVISLIFEKAVFEPTFCPMYAQLCSDLNEKLPSFPSEEPGGKEITFKRVLLNNCQEAFEGAESLRAEIAPRDGEKGQR
KDCQTKNTWKYPSNW*

**>eIFiso4G1-KO8**
MEKDHQPVISLRPGGGGGGPRPGRLFSPAFAAAASGSGDLLRSHVGGASKIGDPNFEVRERVRYTRDQLLELREIVDIPE
AILRINQEIDIELHGEDQIWGRPESDVQVQTQTQAQPHNRYGETDNRDWRARTVQPPAANEEKSWDNIREAKAAHASSGR
QQEQVNRQDQLNHQFASKAQVGPTPALIKAEVPWSARRGNLSEKDRVLKTVKGILNKLTPEKFDLLKGQLMESGITTADI
LKDVISLIFEKAVFEPTFCPMYAQLCSDLNEKLPSFPSEEPGGKEITFKRVLLNNCQEAFEGAESLRAEIAKFDWP*

**>eIFiso4G1-KO11**
MEKDHQPVISLRPGGGGGGPRPGRLFSPAFAAAASGSGDLLRSHVGGASKIGDPNFEVRERVRYTRDQLLELREIVDIPE
AILRINQEIDIELHGEDQIWGRPESDVQVQTQTQAQPHNRYGETDNRDWRARTVQPPAANEEKSWDNIREAKAAHASSGR
QQEQVNRQDQLNHQFASKAQVGPTPALIKAEVPWSARRGNLSEKDRVLKTVKGILNKLTPEKFDLLKGQLMESGITTADI
LKDVISLIFEKAVFEPTFCPMYAQLCSDLNEKLPSFPSEEPGGKEITFKRVLLNNCQEAFEGAESLRAEIAKLTGP*

**>eIFiso4G1-del1**
MEKDHQPVISLRPGGGGGGPRPGRLFSPAFAAAASGSGDLLRSHVGGASKIGDPNFEVRERVRYTRDQLLELREIVDIPE
AILRINQEIDIELHGEDQIWGRPESDVQVQTQTQAQPHNRYGETDNRDWRARTVQPPAANEEKSWDNIREAKAAHASSGR
QQEQVNRQDQLNHQFASKAQVGPTPALIKAEVPWSARRGNLSEKDRVLKTVKGILNKLTPEKFDLLKGQLMESGITTADI
LKDVISLIFEKAVFEPTFCPMYAQLCSDLNEKLPSFPSEEPGGKEITFKRVLLNNCQEAFEGAESLRAEIV--TGPDQEM
ERRDKERIVKLRTLGNIRLIGELLKQKMVPEKIVHHIVQELLGSGPDKKACPEEENVEAICQFFNTIGKQLDENPKSRRI
NDTYFIQMKELTTNLQLAPRLRFMVRDVVDLRSNNWVPRREEIKAKTISEIHDEAMKTLGLRPGATGLTRNGRNAPGGPL
SPGGFPMNRPGTGGMMPGMPGTPGMPGSRKMPGMPGLDNDNWEVPRSKSMPRGDSLRNQGPLLNKPSSINKPSSINSRLL
PHGSGALIGKSALLGSGGPPSRPSSLMASLTHTPAQTAPSPKPVSAAPAVVPVTDKAAGSSHEMPAAVQKKTVSLLEEYF
GIRILDEAQQCIEELQCPEYYSEIVKEAINLALDKGPNFIDPLVRLLEHLHAKKIFKTEDLKTGCLLYAALLEDIGIDLP
LAPALFGEVVARLSLSCGLSFEVVEEILKAVEDTYFRKGIFDAVMKTMGGNSSGQAILSSHAVVIDACNKLLK*

**>eIFiso4G1-del2**
MEKDHQPVISLRPGGGGGGPRPGRLFSPAFAAAASGSGDLLRSHVGGASKIGDPNFEVRERVRYTRDQLLELREIVDIPE
AILRINQEIDIELHGEDQIWGRPESDVQVQTQTQAQPHNRYGETDNRDWRARTVQPPAANEEKSWDNIREAKAAHASSGR
QQEQVNRQDQLNHQFASKAQVGPTPALIKAEVPWSARRGNLSEKDRVLKTVKGILNKLTPEKFDLLKGQLMESGITTADI
LKDVISLIFEKAVFEPTFCPMYAQLCSDLNEKLPSFPSEEPGGKEITFKRVLLNNCQEAFEGAESLRAEIAM-TGPDQEM
ERRDKERIVKLRTLGNIRLIGELLKQKMVPEKIVHHIVQELLGSGPDKKACPEEENVEAICQFFNTIGKQLDENPKSRRI
NDTYFIQMKELTTNLQLAPRLRFMVRDVVDLRSNNWVPRREEIKAKTISEIHDEAMKTLGLRPGATGLTRNGRNAPGGPL
SPGGFPMNRPGTGGMMPGMPGTPGMPGSRKMPGMPGLDNDNWEVPRSKSMPRGDSLRNQGPLLNKPSSINKPSSINSRLL
PHGSGALIGKSALLGSGGPPSRPSSLMASLTHTPAQTAPSPKPVSAAPAVVPVTDKAAGSSHEMPAAVQKKTVSLLEEYF
GIRILDEAQQCIEELQCPEYYSEIVKEAINLALDKGPNFIDPLVRLLEHLHAKKIFKTEDLKTGCLLYAALLEDIGIDLP
LAPALFGEVVARLSLSCGLSFEVVEEILKAVEDTYFRKGIFDAVMKTMGGNSSGQAILSSHAVVIDACNKLLK*

**>eIFiso4G1-del3**
MEKDHQPVISLRPGGGGGGPRPGRLFSPAFAAAASGSGDLLRSHVGGASKIGDPNFEVRERVRYTRDQLLELREIVDIPE
AILRINQEIDIELHGEDQIWGRPESDVQVQTQTQAQPHNRYGETDNRDWRARTVQPPAANEEKSWDNIREAKAAHASSGR
QQEQVNRQDQLNHQFASKAQVGPTPALIKAEVPWSARRGNLSEKDRVLKTVKGILNKLTPEKFDLLKGQLMESGITTADI
LKDVISLIFEKAVFEPTFCPMYAQLCSDLNEKLPSFPSEEPGGKEITFKRVLLNNCQEAFEGAESLRAEIAK-TGPDQEM
ERRDKERIVKLRTLGNIRLIGELLKQKMVPEKIVHHIVQELLGSGPDKKACPEEENVEAICQFFNTIGKQLDENPKSRRI
NDTYFIQMKELTTNLQLAPRLRFMVRDVVDLRSNNWVPRREEIKAKTISEIHDEAMKTLGLRPGATGLTRNGRNAPGGPL
SPGGFPMNRPGTGGMMPGMPGTPGMPGSRKMPGMPGLDNDNWEVPRSKSMPRGDSLRNQGPLLNKPSSINKPSSINSRLL
PHGSGALIGKSALLGSGGPPSRPSSLMASLTHTPAQTAPSPKPVSAAPAVVPVTDKAAGSSHEMPAAVQKKTVSLLEEYF
GIRILDEAQQCIEELQCPEYYSEIVKEAINLALDKGPNFIDPLVRLLEHLHAKKIFKTEDLKTGCLLYAALLEDIGIDLP
LAPALFGEVVARLSLSCGLSFEVVEEILKAVEDTYFRKGIFDAVMKTMGGNSSGQAILSSHAVVIDACNKLLK*

**>eIFiso4G1-del4**
MEKDHQPVISLRPGGGGGGPRPGRLFSPAFAAAASGSGDLLRSHVGGASKIGDPNFEVRERVRYTRDQLLELREIVDIPE
AILRINQEIDIELHGEDQIWGRPESDVQVQTQTQAQPHNRYGETDNRDWRARTVQPPAANEEKSWDNIREAKAAHASSGR
QQEQVNRQDQLNHQFASKAQVGPTPALIKAEVPWSARRGNLSEKDRVLKTVKGILNKLTPEKFDLLKGQLMESGITTADI
LKDVISLIFEKAVFEPTFCPMYAQLCSDLNEKLPSFPSEEPGGKEITFKRVLLNNCQEAFEGAESLRAEIAKLTGPDKE-
------RIVKLRTLGNIRLIGELLKQKMVPEKIVHHIVQELLGSGPDKKACPEEENVEAICQFFNTIGKQLDENPKSRRI
NDTYFIQMKELTTNLQLAPRLRFMVRDVVDLRSNNWVPRREEIKAKTISEIHDEAMKTLGLRPGATGLTRNGRNAPGGPL
SPGGFPMNRPGTGGMMPGMPGTPGMPGSRKMPGMPGLDNDNWEVPRSKSMPRGDSLRNQGPLLNKPSSINKPSSINSRLL
PHGSGALIGKSALLGSGGPPSRPSSLMASLTHTPAQTAPSPKPVSAAPAVVPVTDKAAGSSHEMPAAVQKKTVSLLEEYF
GIRILDEAQQCIEELQCPEYYSEIVKEAINLALDKGPNFIDPLVRLLEHLHAKKIFKTEDLKTGCLLYAALLEDIGIDLP
LAPALFGEVVARLSLSCGLSFEVVEEILKAVEDTYFRKGIFDAVMKTMGGNSSGQAILSSHAVVIDACNKLLK*

**>eIFiso4G1-del6**
MEKDHQPVISLRPGGGGGGPRPGRLFSPAFAAAASGSGDLLRSHVGGASKIGDPNFEVRERVRYTRDQLLELREIVDIPE
AILRINQEIDIELHGEDQIWGRPESDVQVQTQTQAQPHNRYGETDNRDWRARTVQPPAANEEKSWDNIREAKAAHASSGR
QQEQVNRQDQLNHQFASKAQVGPTPALIKAEVPWSARRGNLSEKDRVLKTVKGILNKLTPEKFDLLKGQLMESGITTADI
LKDVISLIFEKAVFEPTFCPMYAQLCSDLNEKLPSFPSEEPGGKEITFKRVLLNNCQEAFEGAESLRAEIAKLTGPDQE-
-------IVKLRTLGNIRLIGELLKQKMVPEKIVHHIVQELLGSGPDKKACPEEENVEAICQFFNTIGKQLDENPKSRRI
NDTYFIQMKELTTNLQLAPRLRFMVRDVVDLRSNNWVPRREEIKAKTISEIHDEAMKTLGLRPGATGLTRNGRNAPGGPL
SPGGFPMNRPGTGGMMPGMPGTPGMPGSRKMPGMPGLDNDNWEVPRSKSMPRGDSLRNQGPLLNKPSSINKPSSINSRLL
PHGSGALIGKSALLGSGGPPSRPSSLMASLTHTPAQTAPSPKPVSAAPAVVPVTDKAAGSSHEMPAAVQKKTVSLLEEYF
GIRILDEAQQCIEELQCPEYYSEIVKEAINLALDKGPNFIDPLVRLLEHLHAKKIFKTEDLKTGCLLYAALLEDIGIDLP
LAPALFGEVVARLSLSCGLSFEVVEEILKAVEDTYFRKGIFDAVMKTMGGNSSGQAILSSHAVVIDACNKLLK*

**>eIFiso4G1-del7**
MEKDHQPVISLRPGGGGGGPRPGRLFSPAFAAAASGSGDLLRSHVGGASKIGDPNFEVRERVRYTRDQLLELREIVDIPE
AILRINQEIDIELHGEDQIWGRPESDVQVQTQTQAQPHNRYGETDNRDWRARTVQPPAANEEKSWDNIREAKAAHASSGR
QQEQVNRQDQLNHQFASKAQVGPTPALIKAEVPWSARRGNLSEKDRVLKTVKGILNKLTPEKFDLLKGQLMESGITTADI
LKDVISLIFEKAVFEPTFCPMYAQLCSDLNEKLPSFPSEEPGGKEITFKRVLLNNCQEAFEGAESLRAEIAKLTGPDQEM
---DKERIVKLRTLGNIRLIGELLKQKMVPEKIVHHIVQELLGSGPDKKACPEEENVEAICQFFNTIGKQLDENPKSRRI
NDTYFIQMKELTTNLQLAPRLRFMVRDVVDLRSNNWVPRREEIKAKTISEIHDEAMKTLGLRPGATGLTRNGRNAPGGPL
SPGGFPMNRPGTGGMMPGMPGTPGMPGSRKMPGMPGLDNDNWEVPRSKSMPRGDSLRNQGPLLNKPSSINKPSSINSRLL
PHGSGALIGKSALLGSGGPPSRPSSLMASLTHTPAQTAPSPKPVSAAPAVVPVTDKAAGSSHEMPAAVQKKTVSLLEEYF
GIRILDEAQQCIEELQCPEYYSEIVKEAINLALDKGPNFIDPLVRLLEHLHAKKIFKTEDLKTGCLLYAALLEDIGIDLP
LAPALFGEVVARLSLSCGLSFEVVEEILKAVEDTYFRKGIFDAVMKTMGGNSSGQAILSSHAVVIDACNKLLK*

**>eIFiso4G1-del9**
MEKDHQPVISLRPGGGGGGPRPGRLFSPAFAAAASGSGDLLRSHVGGASKIGDPNFEVRERVRYTRDQLLELREIVDIPE
AILRINQEIDIELHGEDQIWGRPESDVQVQTQTQAQPHNRYGETDNRDWRARTVQPPAANEEKSWDNIREAKAAHASSGR
QQEQVNRQDQLNHQFASKAQVGPTPALIKAEVPWSARRGNLSEKDRVLKTVKGILNKLTPEKFDLLKGQLMESGITTADI
LKDVISLIFEKAVFEPTFCPMYAQLCSDLNEKLPSFPSEEPGGKEITFKRVLLNNCQEAFEGAESLRAEIAKLTGPDQER
--RDKERIVKLRTLGNIRLIGELLKQKMVPEKIVHHIVQELLGSGPDKKACPEEENVEAICQFFNTIGKQLDENPKSRRI
NDTYFIQMKELTTNLQLAPRLRFMVRDVVDLRSNNWVPRREEIKAKTISEIHDEAMKTLGLRPGATGLTRNGRNAPGGPL
SPGGFPMNRPGTGGMMPGMPGTPGMPGSRKMPGMPGLDNDNWEVPRSKSMPRGDSLRNQGPLLNKPSSINKPSSINSRLL
PHGSGALIGKSALLGSGGPPSRPSSLMASLTHTPAQTAPSPKPVSAAPAVVPVTDKAAGSSHEMPAAVQKKTVSLLEEYF
GIRILDEAQQCIEELQCPEYYSEIVKEAINLALDKGPNFIDPLVRLLEHLHAKKIFKTEDLKTGCLLYAALLEDIGIDLP
LAPALFGEVVARLSLSCGLSFEVVEEILKAVEDTYFRKGIFDAVMKTMGGNSSGQAILSSHAVVIDACNKLLK*

**>eIFiso4G2-Kitaake**
MTQADQAVISLRPGGGGGIGGPRAGRLFPFGASTGSLDFLRPRGGASSGFAAKLGDLRFEPLERVRYTRDQLVELHEIID
IPENILKLKQDIDIELHGEDEPWINNDSSVQTQSYNRYAETDNRDWRSRIEQPVQTPAIGGEEKSWDKFREAKESYISSG
KQDQFNNQDKLSSQFSAKAQVGPAPALVKAEVPWSIQRGNLSNKERVLKTVKGILNKLTPEKFDLLKGQLIEAGITTADI
LKDVISLIFEKAVLEPTFCPMYAQLCFDLNEKLPSFPSEEPGGKEITFKRVLLNNCQEAFEGADNLRSEVNKLTGLDQEM
ERRDKERLVKLRTLGNIRLVGELLKQKMVPEKIVHHIVQELLGSESNRCPAEENVEAICQFFNTIGKQLDENPKSRRFND
VYFNRLKDLTTNSQLASRLRFMARDVLDLRSNQWVPRREEMKAKKISEIHREAENNLGLRPGSTASIRTGRTGTGGGGPL
SPGAFSMNQPGIVGMLPGMPGARKMPGMPGLGSDDWEVPHSRSKPRADPVRNLTPSLANKPSPNNSRLLPQGSAALISGK
TSALVGSGGPLSHGLVVTPSQTTGPPKSLIPAPSVDPIVEQPAAAPKPSSTELQKKTISLLKEYFHILLLHEAQQCIEEL
KSPDYYPEVVKEAINLALDKGTNSIDPLLRLLEHLYNKNVFKATDLETGCLLYSSLLDELAIDLPKAPVHFGEVIGRLVL
SHCLSIEVVEDTLKKIEDSFFRAAVFEAMMKIMKANPSGQAILGSHVAKIDACSKLLSSE*

**>eIFiso4G2-KO1**
MTQADQAVISLRPGGGGGIGGPRAGRLFPFGASTGSLDFLRPRGGASSGFAAKLGDLRFEPLERVRYTRDQLVELHEIID
IPENILKLKQDIDIELHGEDEPWINNDSSVQTQSYNRYAETDNRDWRSRIEQPVSNSCYWW*

**>eIFiso4G2-KO2**
MTQADQAVISLRPGGGGGIGGPRAGRLFPFGASTGSLDFLRPRGGASSGFAAKLGDLRFEPLERVRYTRDQLVELHEIID
IPENILKLKQDIDIELHGEDEPWINNDSSVQTQSYNR*

**>eIFiso4G2-KO3**
MTQADQAVISLRPGGGGGIGGPRAGRLFPFGASTGSLDFLRPRGGASSGFAAKLGDLRFEPLERVRYTRDQLVELHEIID
IPENILKLKQDIDIELHGEDEPWINNDSSVQTQSYNRYKLLLLVVKRNLGTNFVKQKNHTFQVESKINLIIKTN*

**Sup. Table 1 - Number and genotypes of T0 and T2 plants obtained with the different constructs.**

| | OseIFiso4G1 | | | OseIFiso4G2 |
|---|---|---|---|---|
| | p-iso4G1.1 | p-iso4G1.2 | p-iso4G1.3 | p-iso4G2 |
| T0 transgenic plants | 45 | 27 | 18 | 4 |
| T0 edited plants | 37 | 16 | 13 | 2 |
| Independent T0 with mutations leading to | | | | |
| premature stop codon | 11 | 5 | 1 | 3 |
| substitution and/or in-frame deletion | 5 | 1 | 5 | 0 |
| T2 selected for further characterization | | | | |
| with KO mutations | 2 | 1 | 0 | 3 |
| with substitution and/or in-frame deletion | 3 | 0 | 4 | 0 |

**Sup. Table 2 - Segregation of KO and WT alleles in F2 populations derived from crosses between eIFiso4G1 KO and eIFiso4G2 KO lines.** The number of plants observed and the confidence interval (CI) of the percentage (binomial test, p=0,95) is given for each genotype. For comparison, the expected percentage is indicated into brackets. A total of 434 plants have been analysed and derived from crosses iso4G1-KO3 x iso4G2-KO2 (82 plants), iso4G1-KO11 x iso4G2-KO1 (88 plants), iso4G2-KO2 x iso4G1-KO11 (88 plants), iso4G2-KO3 x iso4G1-KO8 (88 plants), iso4G1-KO3 x iso4G2-KO3 (88 plants).

| **Gentoype on *OseIFiso4G2*** | **Gentoype on *OseIFiso4G1*** | | | |
|---|---|---|---|---|
| | **homozygous WT** | **heterozygous** | **homozygous KO** | **all** |
| homozygous WT | 53 plants<br>CI: 9-16 %<br>(6,25 % expected) | 77 plants<br>CI: 14-22 %<br>(12,5 % expected) | 10 plants<br>CI: 1-4 %<br>(6,25 % expected) | 140 plants<br>CI: 28-37 %<br>(25 % expected) |
| heterozygous | 87 plants<br>CI: 16-24 %<br>(12,5 % expected) | 120 plants<br>CI: 23-32 %<br>(25 % expected) | 11 plants<br>CI: 1-5 %<br>(12,5 % expected) | 218 plants<br>CI: 45-55 %<br>(50 % expected) |
| homozygous KO | 42 plants<br>CI: 7-13 %<br>(6,25 % expected) | 34 plants<br>CI: 5-11 %<br>(12,5 % expected) | 0 plants<br>CI: 0-1 %<br>(6,25 % expected) | 76 plants<br>CI : 14-21 %<br>(25 % expected) |
| all | 182 plants<br>CI: 37-47 %<br>(25 % expected) | 231 plants<br>CI : 48-58%<br>(50 % expected) | 21 plants<br>CI: 3-7 %<br>(25 % expected) | 434 plants |

**Sup. Table 3 - Grain filling rate in F2 plants and germination rate in F3 progenies derived from crosses between eIFiso4G1 KO and eIFiso4G2 KO lines.** The grain filling rate was measured on F2 plants homozygous for a KO mutation in one *OseIFiso4G* copy and heterozygous for the other and compared to the expected 75% germination rate if the absence of double mutants were due to female sterility or embryo abortion. The germination rate of F3 progenies was compared to the expected 75% rate if the absence of double mutants were due to a germination defect. Significant differences are indicated by "*" (one-sided binomial test, p<0.05).

| **Cross** | **F2 mother plant** | | | **F3 progeny** | |
|---|---|---|---|---|---|
| | **Genotype on *OseIFiso4G1*** | **Genotype on *OseIFiso4G2*** | **Grain filling rate (Filled grains / Total grains counted)** | **Germination rate** | **Segregation** |
| iso4G1-KO3 x iso4G2-KO3 | KO3/WT | KO3/KO3 | 0,92* (166/181) | 0,71 (89/125) | 77 iso4G1-htz / iso4G2-KO<br>60 iso4G1-KO / iso4G2-KO<br>0 iso4G1-KO / iso4G2-KO |
| iso4G1-KO3 x iso4G2-KO3 | KO3/KO3 | KO3/WT | 0,93* (231/249) | 0,9* 113/125) | - |
| iso4G1-KO11 x iso4G2-KO2 | KO11/WT | KO2/KO2 | 0,88* (287/324) | 0,88* (67/76) | - |
| iso4G1-KO11 x iso4G2-KO2 | KO11/KO11 | KO2/WT | 0,86* (149/174) | 0,82 (51/62) | - |
| iso4G1-KO3 x iso4G2-KO2 | KO3/WT | KO2/KO2 | 0,82 (90/110) | 0,94* (77/82) | - |
| iso4G1-KO3 x iso4G2-KO2 | KO3/KO3 | KO2/WT | 0,86* (89/103) | 0,47 (28/59) | - |

**Sup. Table 4 - Resistance-breaking rates on OseIFiso4G1 KO lines.** Resistance-breaking rates was estimated by the number of plants detected positive with ELISA on leaf samples collected 8 weeks after inoculation with RYMV isolates Ng106, Ng119 or Tg247. Within plant sets inoculated with the same isolate, same letters indicated non significantly different proportions, estimated based on a Fisher exact test.

| Lines | Ng106 | Ng119 | Tg247 |
|---|---|---|---|
| iso4G1-KO3 | 0/60[a] | 0/85[a] | 0/96[a] |
| iso4G1-KO8 | 0/60[a] | 0/71[a] | 0/96[a] |
| iso4G1-KO1 | 0/60[a] | 0/76[a] | 0/96[a] |
| *O. sativa* Gigante (*rymv1.2*) | 2/59[a] | 64/81[b] | 0/96[a] |
| *O. glaberrima* Tog5681 (*rymv1.3*) | 31/61[b] | 67/78[b] | 20/91[b] |

**Sup. Table 5 - Residues involved in key salt-bridges within the VPg central helix–MIF4G hairpin complex.** Color intensity reflects the number of models in which a salt-bridge was predicted between each MIF4G residue and the corresponding VPg residue listed in the first column. MIF4G residue numbering refers to the full-length OseIFiso4G1 protein.

| VPg residues | MIF4G residues | | | |
|---|---|---|---|---|
| | Kitaake WT | iso4G1-del4 | iso4G1-del6 | iso4G1-del7 |
| 38 | 324 | | 317 | |
| 46 | 250 | 243<br>250 | 250 | 243 |
| 48 | 309 | 298<br>309 | 309 | 298<br>309 |
| 50 | | 243 | | 243 |
| 52 | 309 | 298 | 309 | 298<br>309 |
| 56 | 301 | 298<br>301 | 301 | 298<br>301 |

**Sup. Table 6 - Summary of phenotypic traits in the edited lines.** Resistance was primarily assessed with the BF1 isolate and resistance to Cia, Tz11, Mg1 and Ng106 isolates was only assessed on lines resistant to BF1, and the control. Resistance breaking by Ng106, Ng119 and Tg247 was tested for KO lines in *OseIFiso4G1*, using varieties Gigante and Tog5681 carrying resistance alleles *rymv1-2* and *rymv1-3* as controls. Developmental traits were assessed relative to the wild type.

| Lines | Resistance | | | Resistance-breaking | | | Developmental traits | |
|---|---|---|---|---|---|---|---|---|
| | BF1 | Cia, Tz11 | Mg1, Ng106 | Ng106 | Ng119 | Tg247 | plant height | tillering and panicle weight |
| WT | no | no | no | - | - | - | - | - |
| iso4G1-KO3 | high | high | high | not detected | not detected | not detected | mild | not detected |
| iso4G1-KO8 | high | high | high | not detected | not detected | not detected | mild | not detected |
| iso4G1-KO11 | high | high | high | not detected | not detected | not detected | mild | not detected |
| iso4G1-del1 | no | - | - | - | - | - | not detected | not detected |
| iso4G1-del2 | no | - | - | - | - | - | not detected | not detected |
| iso4G1-del3 | no | - | - | - | - | - | not detected | not detected |
| iso4G1-del4 | high | high | high | - | - | - | not detected | not detected |
| iso4G1-del6 | high | high | high | - | - | - | not detected | not detected |
| iso4G1-del7 | partial | partial | no | - | - | - | not detected | not detected |
| iso4G1-del9 | no | - | - | - | - | - | mild | not detected |
| iso4G2-KO1 | no | - | - | - | - | - | mild | not detected |
| iso4G2-KO2 | no | - | - | - | - | - | moderate | not detected |
| iso4G2-KO3 | no | - | - | - | - | - | moderate | not detected |
| Tog5681 | high[1] | high[1] | high[1] | frequent | frequent | frequent | - | - |
| Gigante | high[1] | high[1] | high[1] | rare | frequent | not detected | - | - |

'-' indicates phenotypic traits that have not been assessed

[1]according to Hébrard E, Pinel-Galzi AA, Oludare A, Poulicard N, Aribi J, Fabre S, Issaka S, Mariac CC, Dereeper A, Albar L, Silue D, Fargette DJ (2018) Identification of a hypervirulent pathotype of Rice yellow mottle virus : a threat to genetic resistance deployment in West-Central Africa. Phytopathology, 108, 299–307. https://doi.org/10.1094/PHYTO-05-17-0190-R

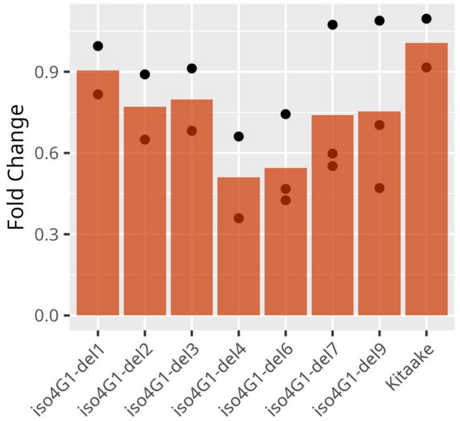


**Sup. Fig. 1. Expression level of *OseIFiso4G1* in lines with deletions in the central domain of the gene.** Relative expression of OseIFiso4G1 was determined by RT-qPCR using actin as reference. Two or three biological replicates were used for each line. No significant difference between lines was detected with a Kruskall-Wallis test (p-value = 0.3).

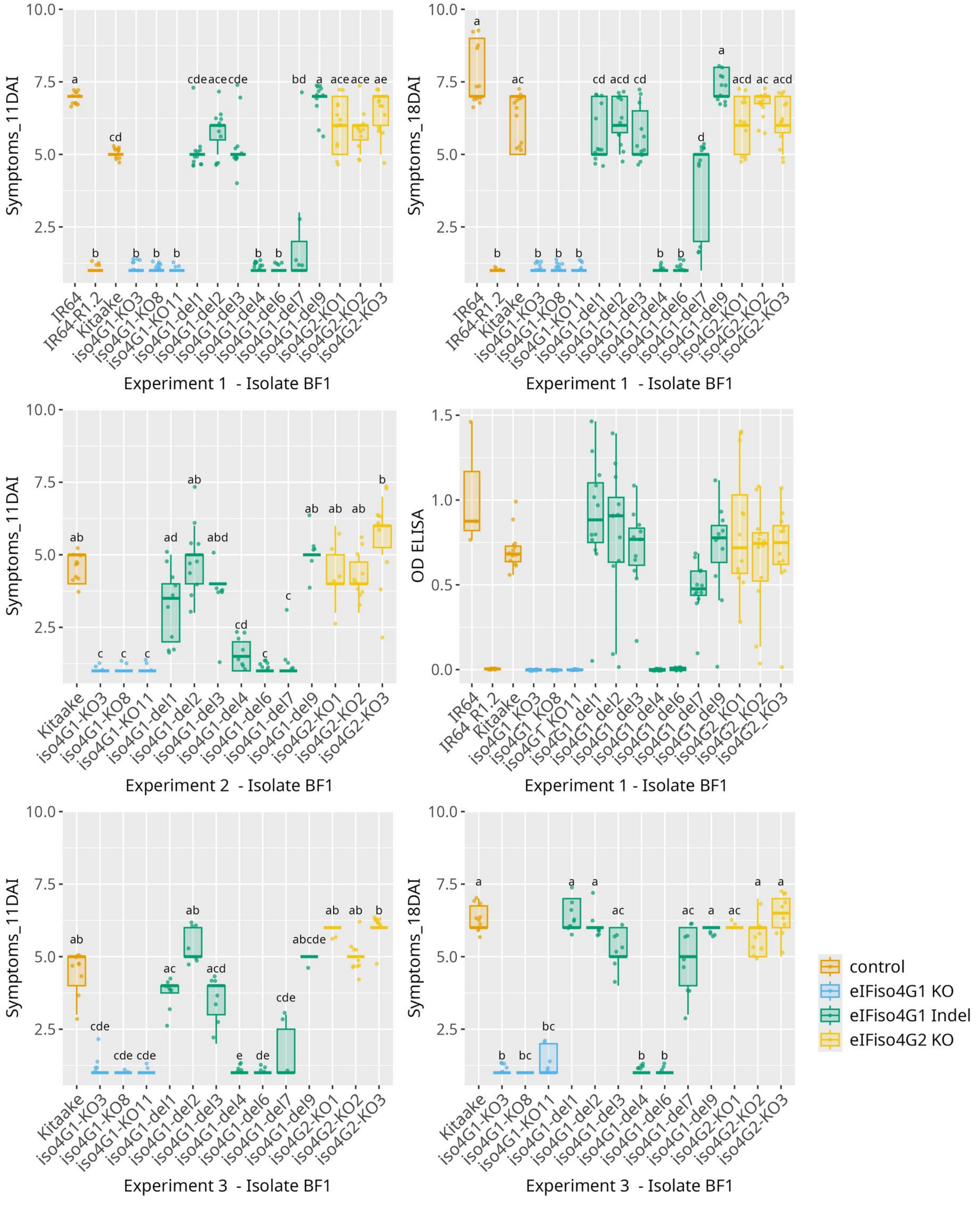


**Sup. Fig. 2. Resistance of edited lines against BF1 isolate of RYMV.**

Resistance was evaluated based on the symptoms observed 11 and 18 DAI and ELISA on leaf samples collected 13 (experiment 2) or 15 (experiment 1) DAI. ELISA was not performed on the third experiment. Kitaake wild-type and IR64 were used as susceptibility control ; IR64 introgressed with rymv1.2 resistance allele (IR64_R1.2) was used as resistant control. The lower and upper hinges correspond to the first and third quartiles ; the upper vs. lower whisker extend from the hinge to the largets vs. smallest value no further than 1,5 x interquartile from the hinge. Different letters above the box-plots represent significant differences based on a Kruskall-Wallis test, followed by a Dunn post-hoc test (p-value <0,5 after fdr correction).

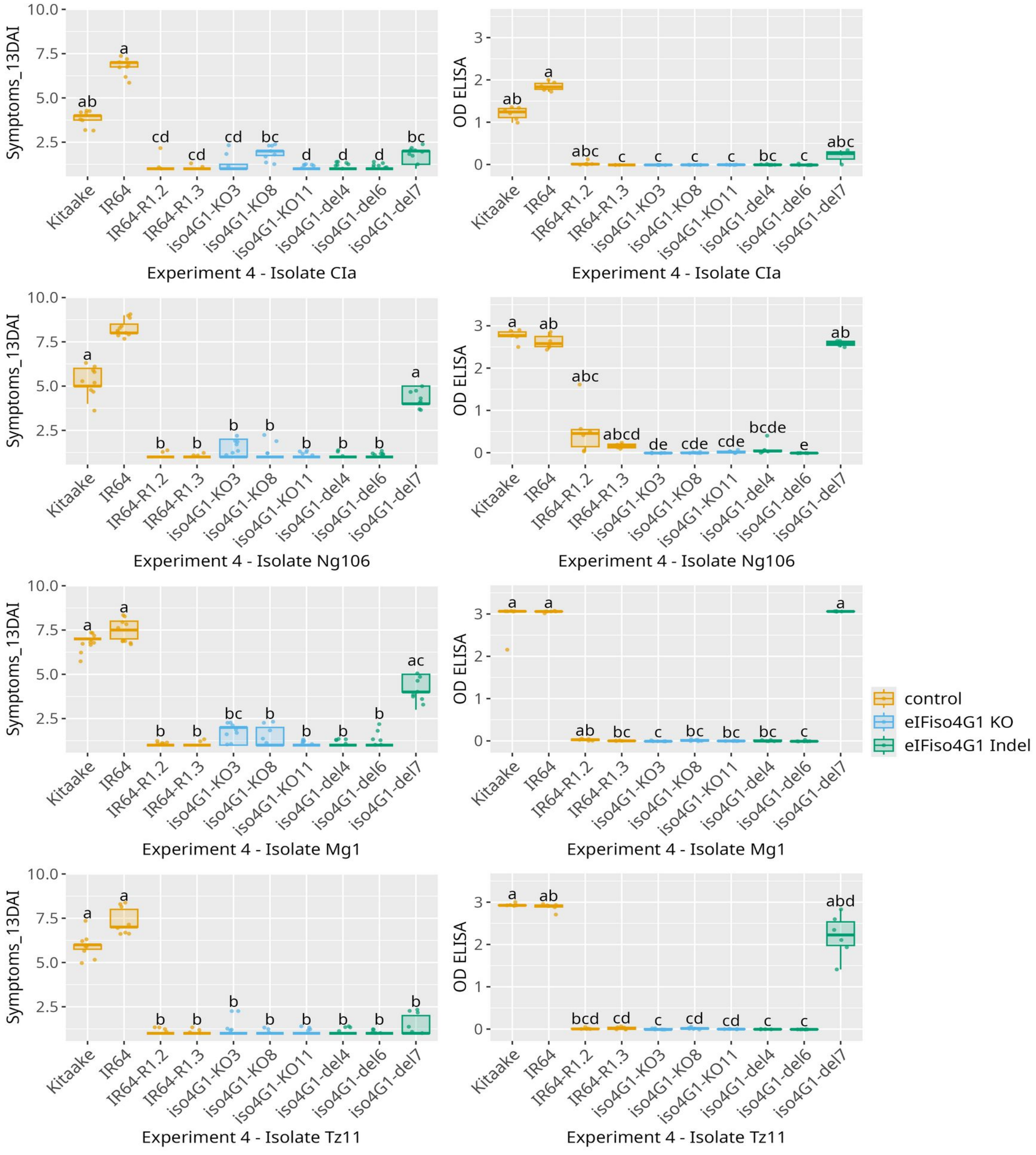


**Sup. Fig. 3 - Resistance of edited lines against isolates CIa, Ng106, Mg1 and Tz11.**
Only lines resistant with isolated BF1 were tested with additional isolates. Symptoms observed 13 DAI and ELISA on leaf samples collected 13 (for Ng106 and Mg1) or 14 (for CIa and Tz11) DAI are represented. Kitaake wild-type and IR64 were used as susceptibility controls ; IR64 introgressed with rymv1.2 (IR64_R1.2) or rymv1.3 (IR64_R1.3) resistance alleles were used as resistant controls. The lower and upper hinges correspond to the first and third quartiles ; the upper vs. lower whisker extend from the hinge to the largets vs. smallest value no further than 1,5 x interquartile from the hinge. Different letters above the box-plots represent significant differences based on a Kruskall-Wallis test, followed by a Dunn post-hoc test (p-value <0,5 after fdr correction).

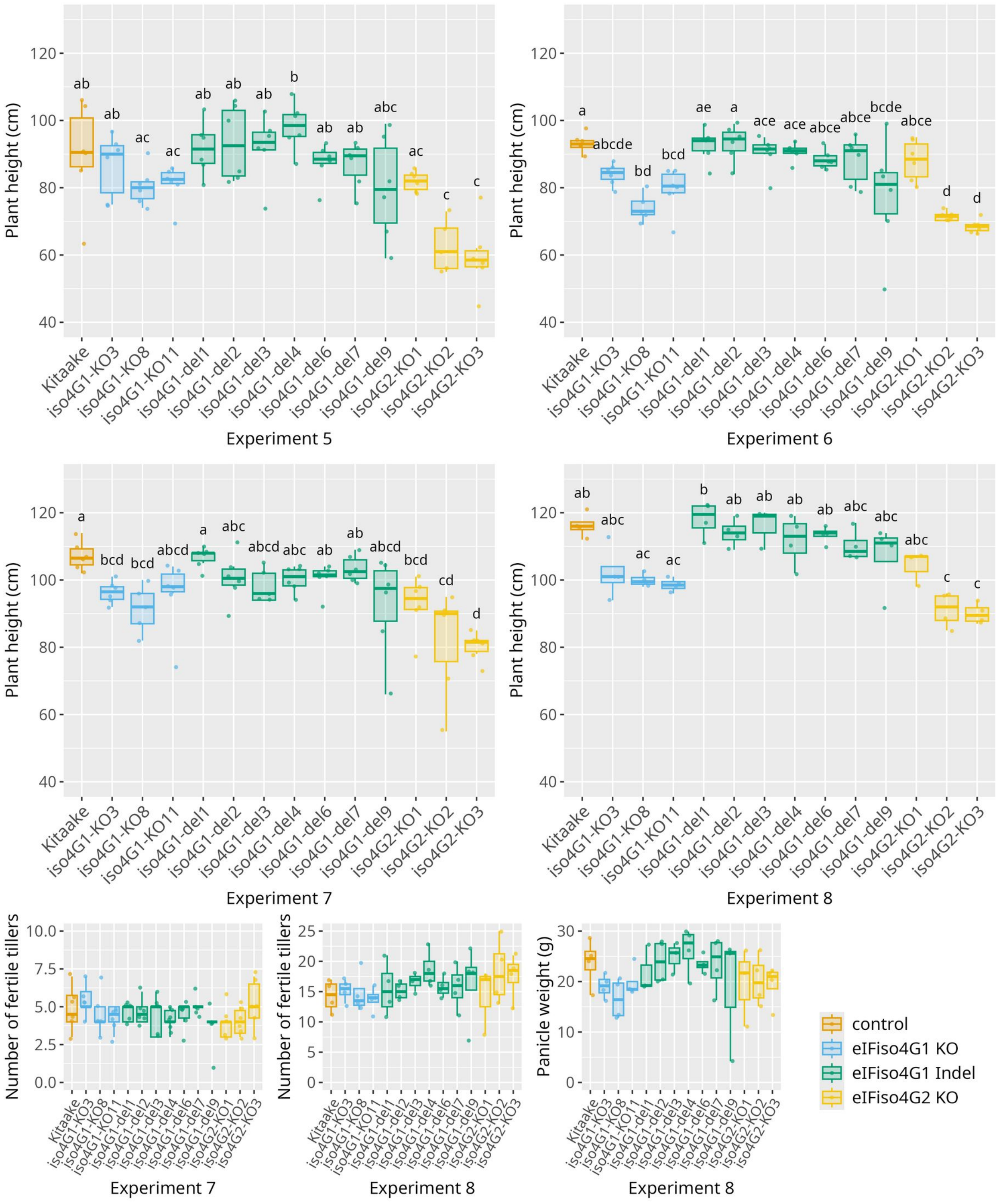


**Sup. Fig. 4 - Morphology of edited lines in controlled conditions.**

Edited lines were evaluated for plant height at ripening stage in four independent experiments performed in controlled conditions. Results of three experiments are represented here, while results of the fourth one is presented in the main document. Fertiles tillers number was measured in two experiments and seed weight in a single one. The lower and upper hinges correspond to the first and third quartiles ; the upper vs. lower whisker extend from the hinge to the largets vs. smallest value no further than 1,5 x interquartile from the hinge. Different letters above the box-plots represent significant differences based on a Kruskall-Wallis test, followed by a Dunn post-hoc test (p-value <0,5 after fdr correction).

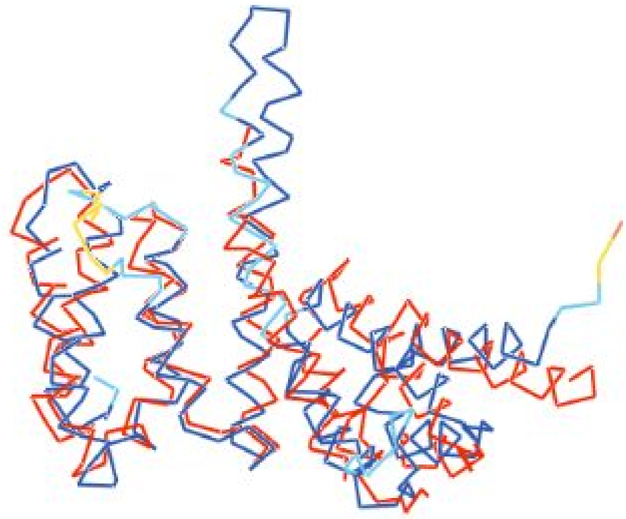

**Sup. Fig. 5 - Comparison between the predicted 3D model of the rice MIF4G domain (colored by plDDT) and the crystallographic structure of human MIF4GII (PDB 1HU3, shown in red), displayed in side view.** The predicted structure of the Kitaake MIF4G domain was superimposed onto the human MIF4GII crystal structure. As expected given the 32% sequence identity between the two proteins, they exhibited strong structural similarity with root mean square deviations (RMSD) below 2.5 Å. Notably, the two central helices 3a and 3b were predicted to form a longer hairpin in rice.

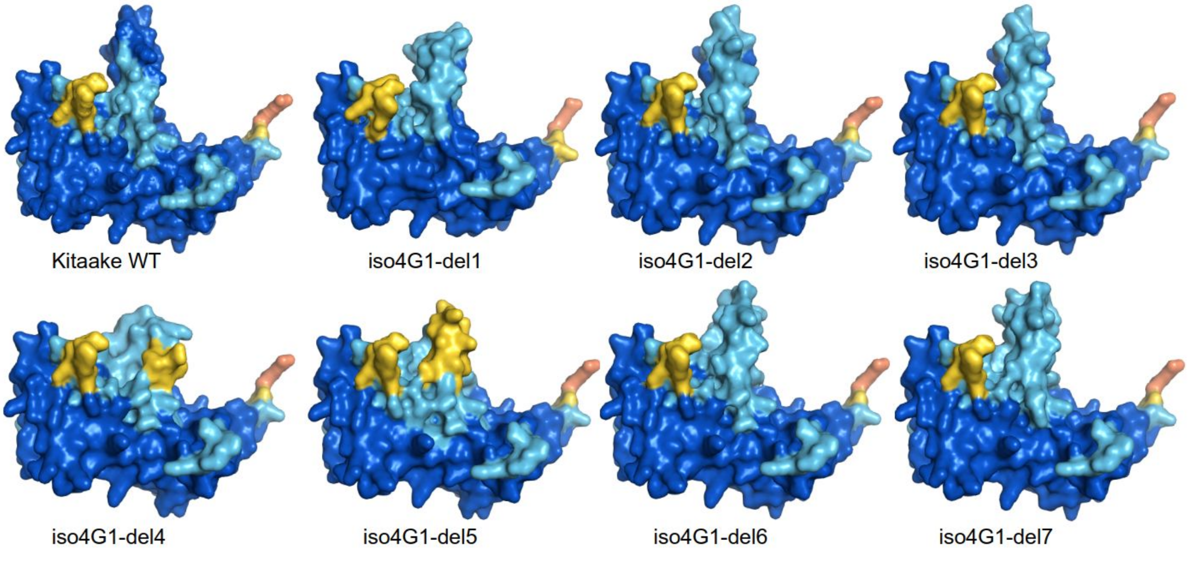


**Sup. Fig. 6 – Predicted MIF4G 3D surface models of WT Kitaake and edited variants.** All models were superimposed in the same orientation for comparison. Structures are colored according to per-residue model confidence score (plDDT): dark blue (plDDT>90) indicates high-confidence regions; light blue (plDDT>80), confident regions; yellow (plDDT>60) and orange (plDDT>50), low-confidence regions.

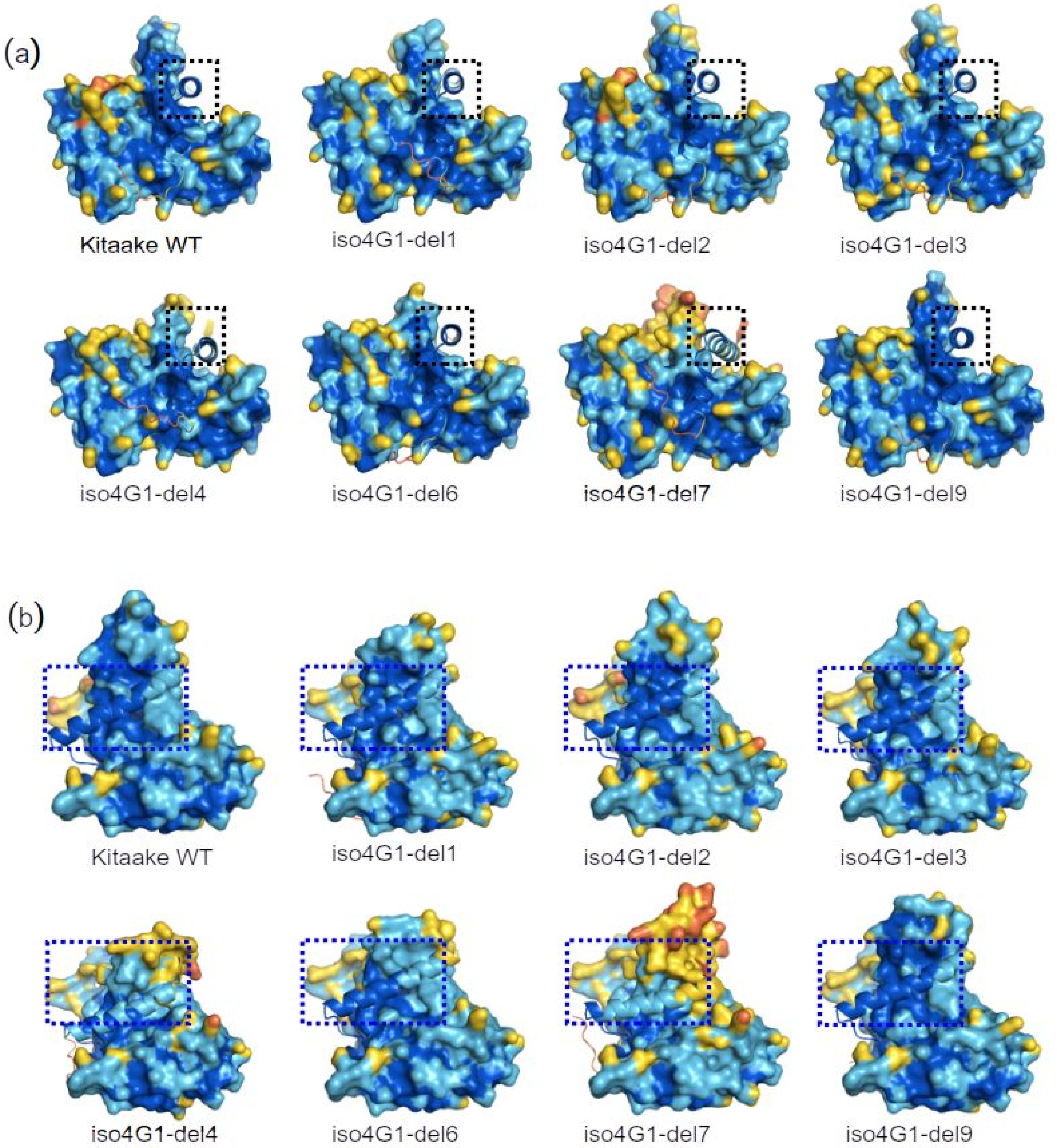


**Sup. Fig. 7 – Predicted 3D structure of MIF4G/VPg complex for WT Kitaake and edited variants, represented in side (a) and front (b) views.** VPg and MIF4G are shown as ribbon and surface representations, respectively. The dotted frames highlight differences in VPg helix length and position among variants. For each view, all complexes were superimposed in the same orientation for comparison. The N-terminal domain of VPg was masked for clarity. Structures are colored according to per-residue model confidence score (plDDT): dark blue (plDDT>90) indicates high-confidence regions; light blue (plDDT>80), confident regions; yellow (plDDT>60) and orange (plDDT>50), low-confidence regions.

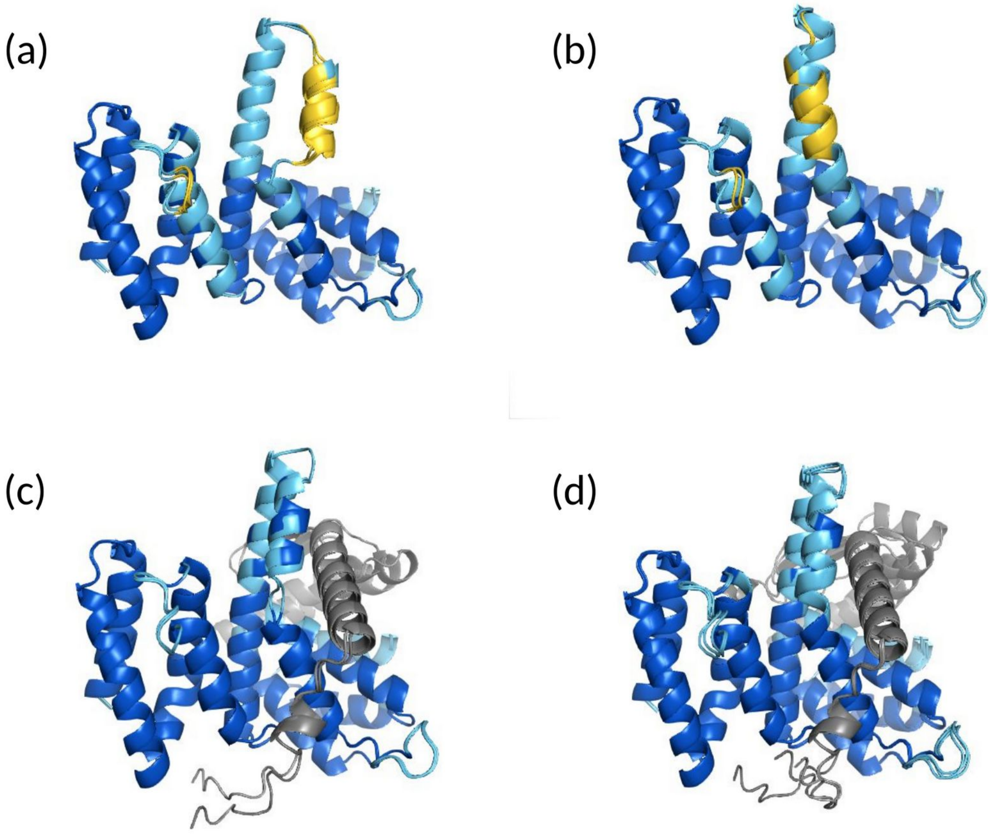


**Sup. Fig. 8 – Conformational variability in the predicted 3D structures of the MIF4G domain for the rymv1-3 allele alone (a, b) or in complex with RYMV VPg (c,d).** Two out of five models showed a broken helix 3a (a, c) while it is not the case in the other three models (b, d). The MIF4G structures are colored according to per-residue model confidence score (plDDT), with the following scheme: dark blue (plDDT > 90) for high-confidence regions, light blue (plDDT > 80) for confident regions, yellow (plDDT > 60) for low-confidence regions, and orange (plDDT > 50) for regions with the lowest confidence. The central helix of VPg is colored grey.
